# Accounting for RNA polymerase heterogeneity reveals state switching and two distinct long-lived backtrack states escaping through cleavage

**DOI:** 10.1101/762468

**Authors:** Richard Janissen, Behrouz Eslami-Mossallam, Irina Artsimovitch, Martin Depken, Nynke H. Dekker

## Abstract

Pausing by bacterial RNA polymerase (RNAp) is vital in the recruitment of regulatory factors, RNA folding, and coupled translation. While backtracking and intra-structural isomerization have been proposed to trigger pausing, our understanding of backtrack-associated pauses and catalytic recovery remains incomplete. Using high-throughput magnetic tweezers, we examined the *E. coli* RNAp transcription dynamics over a wide range of forces and NTP concentrations. Dwell-time analysis and stochastic modeling identified, in addition to a short-lived elemental pause, two distinct long-lived backtrack pause states differing in recovery rates. We further identified two stochastic sources of transcription heterogeneity: alterations in short-pause frequency that underlie elongation-rate switching, and RNA cleavage deficiency that underpins different long-lived backtrack states. Together with effects of force and Gre factors, we demonstrate that recovery from deep backtracks is governed by intrinsic RNA cleavage rather than diffusional Brownian dynamics. We introduce a consensus mechanistic model that unifies our findings with prior models.

## INTRODUCTION

The transcription of genetic information is the first step in gene expression. The synthesis of RNA transcripts is carried out by the DNA-dependent RNA polymerases, whose structures and molecular mechanisms are largely conserved throughout all domains of life (1). Transcription is a complex and tightly controlled process that defines cell identity, fitness, proliferation, and response to environmental signals. The RNA polymerase itself is subjected to many forms of regulation, which include interactions with accessory proteins, nucleic acid signals, and small molecules inducing conformational changes in the polymerase active site and global structural changes in the transcription elongation complex (TEC) (2–5).

Pioneering work on bacterial RNA polymerases (RNAp) revealed that RNA chain elongation is intermittent, and frequently interrupted by pauses that reduce the overall transcription velocity (6, 7). Numerous subsequent studies, reviewed in (2), identified two types of pauses: sequence-induced regulatory pauses that allow timely interaction with transcription factors and the translation machinery, and stochastic pauses originating from intrinsic random events within the enzyme (e.g. structural isomerization, misincorporation), both promoting RNAp backtracking relative to the DNA template and the RNA. Over the past two decades, different high-resolution single-molecule techniques have identified an ubiquitous catalytically inactive elemental pause state, *cis*-acting consensus DNA sequences, and interruption of active RNA chain elongation induced by backtracking in bacterial RNAp (and also in eukaryotic Pol II) (8–13). While short intrinsic pauses with lifetimes about 1 s have been identified as signatures originating from the elemental pause in RNAp, pauses with longer lifetimes comprise a rather heterogeneous class of intermediates (5, 14). Apart from rare RNA hairpin-stabilized pauses, long-lived pauses are commonly attributed to backtracked RNAp. Despite Gre factor-mediated cleavage facilitating recovery from backtracked states, previous biochemical studies also emphasized the role of intrinsic cleavage in recovery from deep backtracks, and proposed that RNAp might undergo two distinct backtrack states based on the observation of two different intrinsic RNA cleavage rates (∼5 s and ∼100 s) (5, 15–17). Analogous to this hypothesis are the observations made for eukaryotic Pol II, which recovers from deeper backtracks predominantly by RNA cleavage rather by diffusional Brownian motion (18). However, to date most single-molecule studies on RNAp have attributed all pauses exceeding 5 s to a single state with a broad power-law distribution of pause durations that originate from diffusive TEC dynamics during RNAp backtracking, and consider thus diffusional Brownian motion of the RNAp to be the main recovery pathway (11, 12, 19–22). Due to these divergent interpretations of transcriptional pause signatures from bulk and single-molecule experiments, a consensus mechanistic model describing intrinsic transcriptional pausing and backtrack recovery for RNAp remains lacking.

Previous single-molecule studies may not have fully captured pause states whose occurrences were infrequent within limited observation times. To test this hypothesis, we employed high-throughput magnetic tweezers to probe the full temporal spectrum of *E. coli* RNAp transcription dynamics over a time period exceeding two hours. Through dwell-time analysis and simulation-based maximum-likelihood parameter estimation – which accounts for colored experimental noise and the effects of trace smoothing – we identified and fully characterized three distinct pause states that compete with elongation: a short-lived pause with a lifetime of ∼1s, which resembles the reported consensus elemental pause, and two long-lived, serially connected pause states, with lifetimes of ∼4 s and ∼100 s, that branch from the elemental pause. Our results further showed that the recovery from backtracks of at least 4 nucleotides is dominated by intrinsic RNA cleavage and not by diffusional Brownian motion, and that a subset of paused TECs assumes a conformation in which intrinsic and GreB-assisted RNA cleavage are hindered, delaying the escape to productive elongation. Our large datasets also allowed us to investigate the poorly understood heterogeneity in transcription velocity and pause dynamics, providing evidence for previously postulated state-switching (14, 23) and demonstrating that it derives from stochastic alterations in the frequency of short pauses. By integrating these key findings with the results of previous studies, we present a unified mechanistic model that fully describes the origin and hierarchy of intrinsic pause states.

## RESULTS

### RNAp transcription dynamics exhibit notable heterogeneity

To be able to capture infrequent events potentially missed by previous investigations, we established a single-molecule assay (**Methods M1**) based on high-throughput magnetic tweezers with instrumental drift compensation (**Methods M2**) that provides large data sets suited to statistically robust analysis (24, 25). Halted TECs were formed on surface-attached linear DNA templates by nucleotide deprivation, as described previously (26), on a template encoding the *E. coli rpoB* gene free of known consensus pause-inducing sequences that could overshadow the intrinsic RNAp pausing dynamics (10, 12). By attaching magnetic beads to singly biotinylated RNAps and varying the DNA template orientation (**Fig. 1A**), we could apply assisting forces (AF) or opposing forces (OF) to the RNAps while monitoring their progression along the DNA template in real time. Transcription was restarted upon the addition of all four nucleotides in equimolar concentration, and monitored for ≥2 hours at constant force.

**Figure 1:**
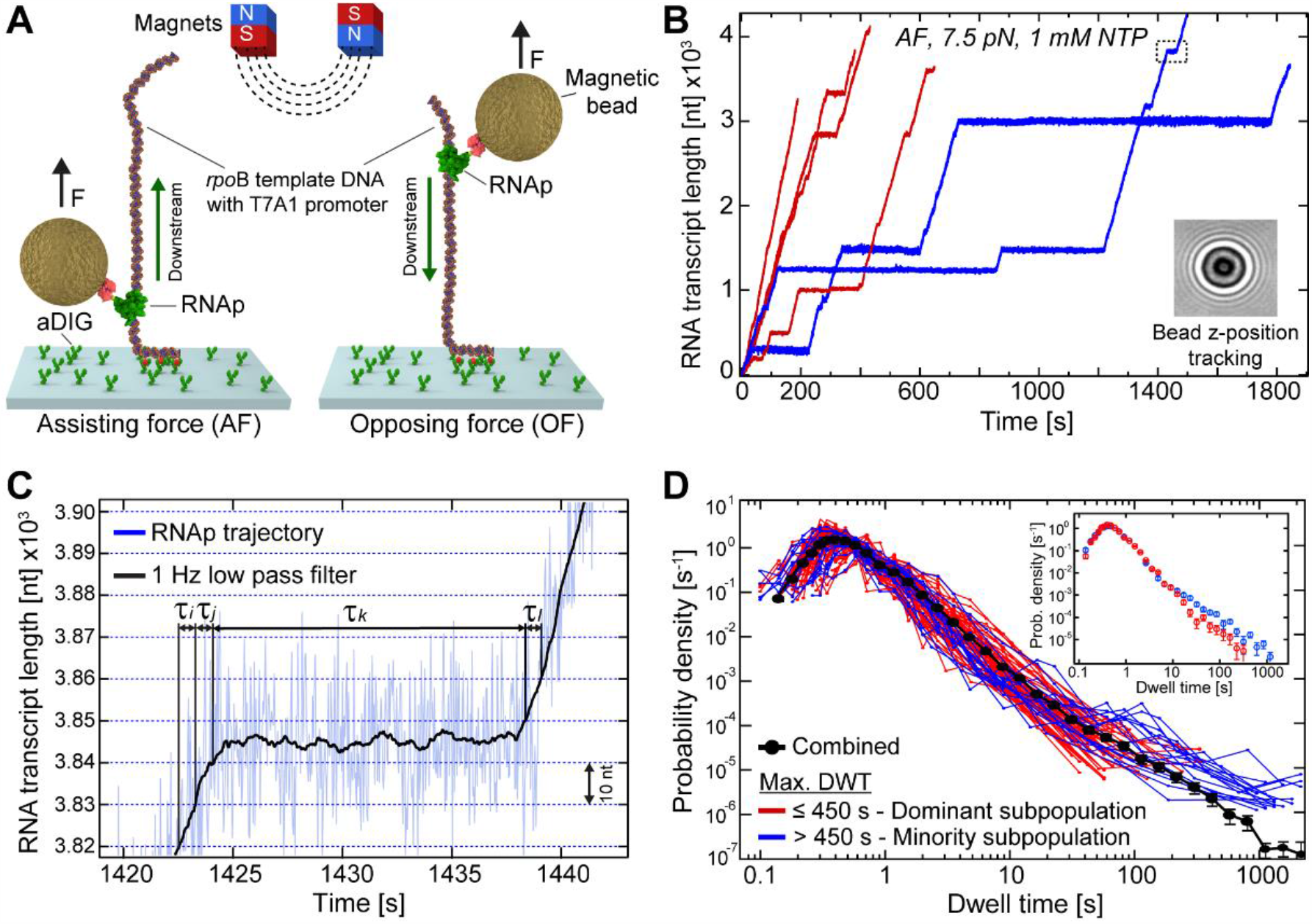
Single-molecule RNAp transcription dynamics reveals notable heterogeneity among individual trajectories. (**A**) Schematic representation of the magnetic tweezers *in vitro* transcription assay in the assisting force (AF) and opposing force (OF) configurations with respect to the RNAp downstream direction. The assay consisted of a surface-attached DNA template including a T7A1 promoter sequence, a magnetic bead bound to the singly biotin-labeled ß’-subunit of RNAp, and the magnets that exerted a constant force *F* on this bead. (**B**) Example trajectories of individual RNAps. Translocation was measured *via* the height change of the bead diffraction pattern (inset) over time. Individual trajectories are colored based on their largest pause duration according to the populations identified in (D). (**C**) Magnified region of an individual RNAp trajectory corresponding to the data in the dashed rectangle in (B). Dwell times (denoted in the example with *τ*_*i*_, *τ*_*j*_, *τ*_*k*_ and *τ*_*l*_) associated with RNAp advancing consecutively through windows of several nucleotides (10 nt windows chosen for facilitated visualization, blue dashed horizontal lines) were extracted from 1 Hz-filtered trajectories (black line). (**D**) Superimposition of DWT distributions from individual trajectories (blue, red) and the combined DWT distribution of all trajectories (black, *n* = 196 individual trajectories). The color code reflects the largest pause duration (maximum DWT) found for each individual RNAp trajectory (red: ≤ 450 s; blue: > 450 s), separated by the established selection protocol (Methods M4). The inset depicts the combined DWT distributions of the two separated subpopulations (blue, red). **See also Figure S1**.

In agreement with previous studies, the RNAp trajectories (**Fig. 1B**) exhibit periods of apparently constant velocities punctuated by pauses of different duration and position (14, 26). We constructed dwell-time (DWT) distributions from extracted DWTs (**Fig. 1C**) obtained for all trajectories (24) by measuring the time needed for each individual RNAp to successively transcribe a fixed number of nucleotides (**Methods M3**). The statistics of analyzed RNAp trajectories, total transcript lengths, and DWTs are specified in **Table S1** for all different conditions tested. With the extracted DWTs, we could then perform a probabilistic analysis of RNAp kinetics sampled from a hundred milliseconds up to two hours and extract the underlying kinetic parameters for RNA chain elongation and transcriptional pausing (24, 27).

**Figure 1D** depicts individual example DWT distributions for trajectories measured under 7.5 pN AF at 1 mM [NTPs]. All individual DWT distributions qualitatively exhibit the same features: a peak at short time scales (∼400 ms for the example shown), and a tail of gradually decreasing probability density for DWT >1 s. While the peak reflects the pause-free elongation, the tail originates from pauses (24, 26). We observe a significant variation between individual DWT distributions, particularly in the occurrence of extremely long-lived pauses. While for the majority of trajectories all pauses lie well below 450 s (**Fig. 1D**, red), for a small subpopulation (∼ 10%) of trajectories the pauses extend to few thousand seconds (**Fig. 1D**, blue). We found that selecting the trajectories for long pauses (>450 s) increases the likelihood to observe not only the longest pauses, but also shorter pauses all the way down to ∼10 s **(Fig. 1D, inset)**, indicating that these trajectories actually belong to a distinct subpopulation of RNAps. As a result, the combined DWT distribution (**Fig. 1D**, black), constructed by accumulating measured DWTs from all single trajectories without any preselection, is not representative of any of the individual DWT distributions (**Fig. 1D**, red and blue). To reveal differences between these subpopulations, we used the Bayes-Schwarz information criterion (**Fig. S1A**,**B**,**C**) to estimate the number of single-rate pause states in the combined DWT distribution and in the DWT distributions corresponding to each subpopulation (28, 29). We found that while the pausing dynamics of each subpopulation can be captured with three single-rate pause states, the combined DWT distribution requires the incorporation of a fourth pause state, an artefact arising from combining two inherently different distributions. Since we found that the maximum pause duration provides a parameter that can successfully discriminate between the two subpopulations, we established a selection protocol using the Gaussian mixture model approach, which classifies individual trajectories based on the longest pause they contain (**Methods M4**). We first discuss the dominant sub-population, and then return to dissect the minority sub-population together with the verification that the detected subpopulations represent actual difference in RNAp endonuclease activity.

### Intrinsic RNAp dynamics reflects three distinct pause states

Previous studies have shown that the consensus elemental pause (lifetime ∼1 s) can act as an intermediate state for less frequent “stabilized” pauses with extended lifetimes (2, 30). All other intrinsic pause signatures were previously attributed to diffusive backtracking. However, our measured DWT distribution is incompatible with the simple power-law decay that associates diffusive entry to, and recovery from, backtracking (grey dashed line in **Fig. 2A**). In contrast, the Bayes-Schwarz information criterion (BIC) analysis supports the existence of three distinct pause states (**Figs. 2A, S1D, S2A**; **Methods M5, M6**). We therefore propose a working model (**Fig. 2**) that includes a frequent short pause state (P1) and two additional rare pauses (P2, P3) of extended duration. The catalytic pathway is characterized by an effective elongation rate *k*_el_, as RNA chain elongation consists of several sub-processes including nucleotide addition, NTP hydrolysis, PP_i_ release, and translocation. It thus has non-exponential transition times to both pausing and subsequent RNA chain elongation. The effective elongation rate *k*_el_ should be considered as the inverse of the average time it takes to complete one nucleotide-addition cycle, conditioned on RNAp not entering a pause (see also further discussion of *k*_el_ related to **Fig. 3D**). Here, we assume that a typical time associated with entering a pause is much shorter than the time to exit a pause, and ignore it in what follows. Therefore, we only consider the probability of leaving the catalytic pathway and entering the off-pathway (through P1), and denote this by *P*_total_. The three pause states (P1, P2, and P3) are connected sequentially, jointly forming a single off-pathway branch; the entry into P2 and P3 is governed by the rates *k*_P2_ and *k*_P3_, respectively, and all pauses return to the catalytic pathway through the exit rates *k*_e1_, *k*_e2_, and *k*_e3_.

**Figure 2:**
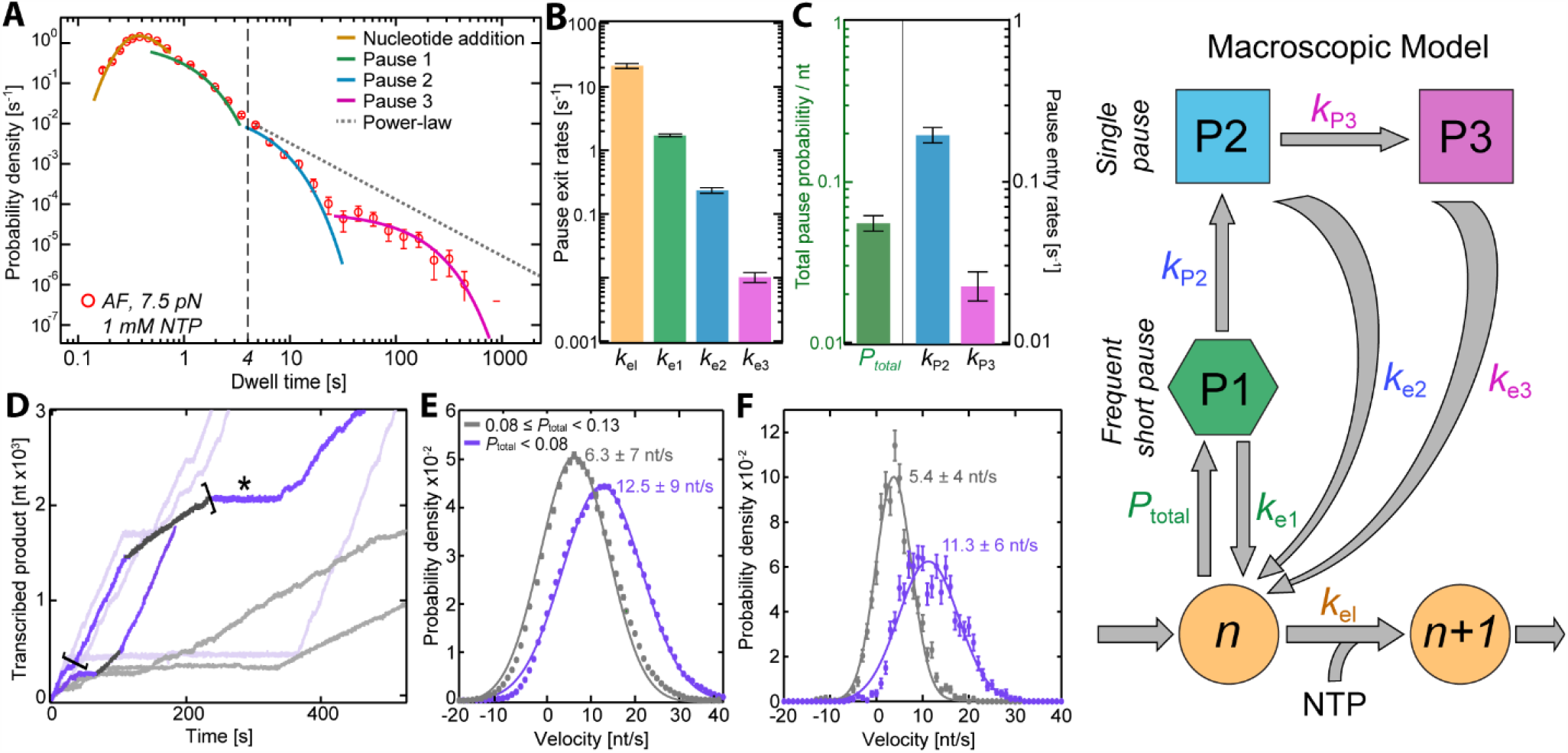
Analysis and extraction of kinetic values of RNAp transcription dynamics. (**A**) DWT distribution (red circles) for *n* = 230 individual trajectories measured under 7.5 pN AF at 1 mM NTPs. Solid curves show the best fit of the model (rightmost panel) to the data set, where the contributions of individual components of the model are separated: the orange curve at short timescales captures the effective pause-free elongation, while the three curves covering the range ∼1 s to ∼1,000 s correspond to three identified exponential pauses (P1: green, P2: cyan, P3: magenta), determined by BIC. The dashed black line depicts the pause lifetime threshold of 4 s. The grey dashed line illustrates a power-law distribution expected from diffusive backtracking dynamics. (**B**) and (**C**) show the values of the model parameters from fitting the model to the DWT distribution in (A). The coloring reflects the different kinetic states in the model. One-sigma confidence intervals are determined by bootstrapping with 1,000 iterations. (**D**) Example trajectories showing stochastic switching of velocity (bold coloring). State switching is not detectable in the majority of trajectories (transparent coloring), although a significant variation in velocities (between purple and grey colored trajectories) is evident. (**E**) Distribution of velocities for the two groups of trajectories, separated by their estimated pause probabilities (*P*_total_). The error denotes the 1-σ confidence interval obtained by bootstrapping 100 times. (**F**) Velocity distribution obtained for two segments of a single trajectory (*) from (D; segments enclosed in brackets). Solid lines and values in (E) and (F) show the best Gaussian fits. **See also Figures S1, S2, and S3**.

**Figure 3:**
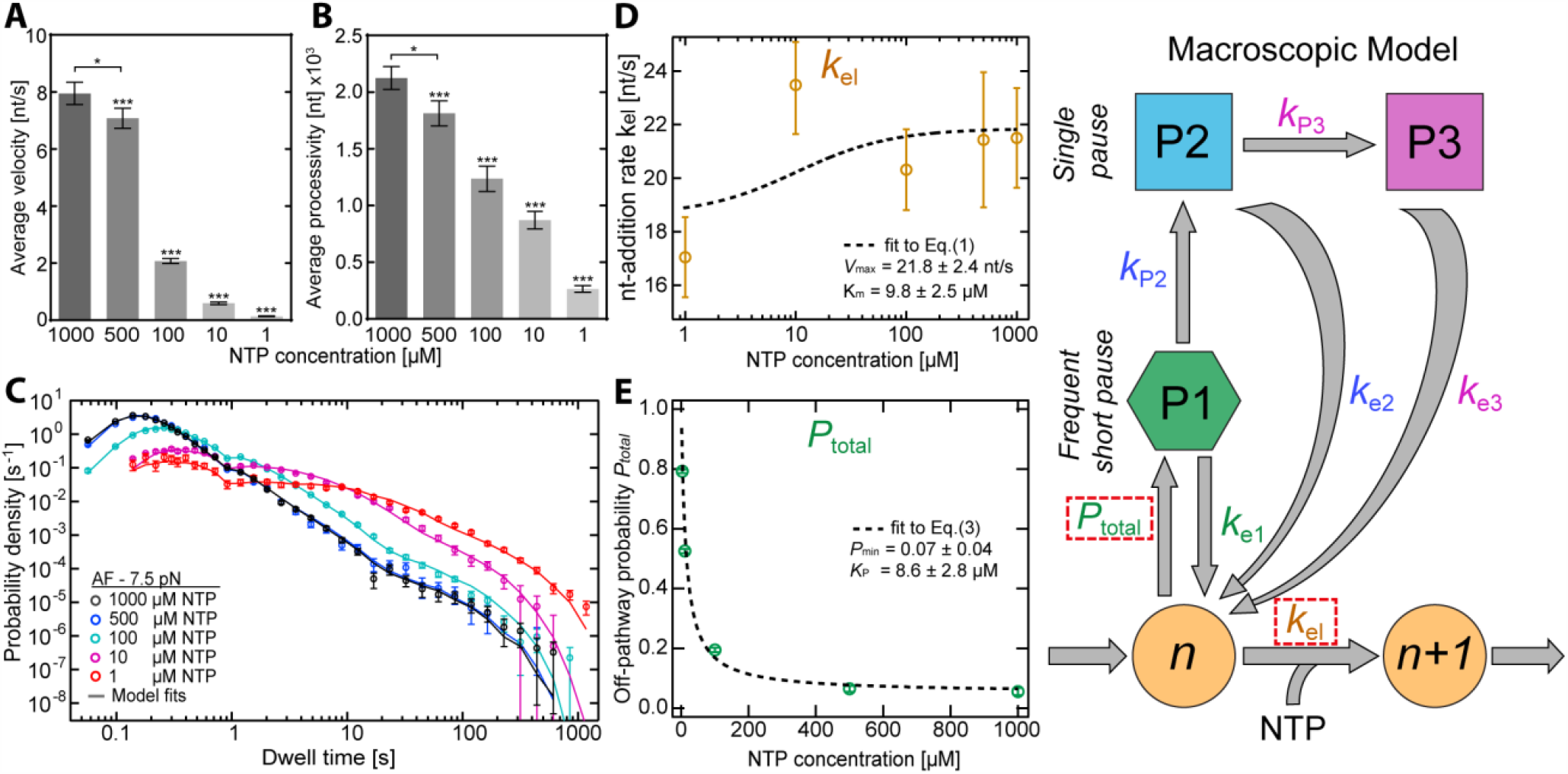
NTP deficiency increases pausing probability and reduces effective elongation rate. (**A**) Measured average (± SEM) end-to-end velocity and (**B**) processivity (± SEM) at different [NTPs]. (**C**) Superimposed DWT distributions (circles) at different [NTPs]. The lines represent the best two-parameter fit of the model (rightmost panel) to each data set by only setting *k*_*el*_ and *P*_total_ as free fit parameters (dashed rectangles). (**D**) Effective *k*_el_ from (C), fitted with **Eq. 1** (dashed line). (**E**) Off-pathway entry probabilities *P*_total_ resulting from the fits to data sets in (C), fitted with **Eq. 3** (dashed line). Data shown in (A, B) were subject to statistical analysis using one-way ANOVA, with comparative Tukey post-hoc test (significance level *α*: *\*\*\* = 0*.*001* and *\* = 0*.*05*). Error bars for (C, D, E) were determined by bootstrapping. **See also Figure S4**.

Though many other pause configurations could explain any one of the data sets, we now proceed to rule these out by demanding that our model is simultaneously consistent with a wide variety of experimental conditions. As a first step, we parametrized the working model by fitting it to a combined DWT distribution, constructed from a large (361.3 kb of total transcript length; **Table S1**) reference ensemble of trajectories measured under 7.5 pN AF at a saturating [NTPs] of 1 mM. Under these conditions, the characteristic pause times (**Fig. 2A**; determined by the green, cyan, and magenta “shoulders”) are well-separated from the pause-free elongation peak (**Fig. 2A**; orange peak), which allows for accurate maximum-likelihood estimation of all parameters (**Methods M6**; **Fig. 2B**,**C**; **Table S2**) (24). The parameter values estimated from the reference data set will be referred to as reference parameters in all subsequent analyses. Having confirmed that the fit remains robust and is independent of the DWT-window size (we evaluated 4, 10 and 20 nt DWT windows; **Fig. S2B, C**), we chose a 4-nt window for all subsequent analyses.

We found that the model captures the velocity distribution (**Fig. S2D, E**; **Methods M7**) used in previous studies to estimate the *k*_el_ (12, 22, 31, 32). The estimated *k*_el_ (**Fig. S2F**) from the velocity distribution (*k*_el_: 21.6 ±0.2 nt/s) is compatible both with the value we obtain from DWT-distribution analysis (*k*_el_: 21.7 ±1.8 nt/s) and with reported pause-free velocity estimations (22, 32). Similarly, the lifetime of ∼1 s for the frequent pause state P1 is similar to that of the elemental pause, and pause lifetimes compatible with our estimates of ∼4 s (P2) and ∼100 s (P3) have also been reported (5, 15, 16, 20, 33).

We next investigate the variation in transcription velocity, which we define as the velocity observed between longer pauses (P2 and P3). To focus on the transcription velocity for individual RNAp, we removed all DWTs >4 s (which encompasses the P2 and P3 lifetimes; dashed line in **Fig. 2A**). We then calculated the displacement of RNAp in consecutive temporal windows of 1 s. We also estimated the values of *k*_el_ and *P*_total_ for individual trajectories in the reference data set (**Fig. S3A**; **Methods M7**), and we found that both quantities follow wide distributions (**Fig. S3A**,**B**,**C**). While the distribution for *k*_el_ is largely symmetric, the distribution for *P*_total_ is skewed and bimodal, and the transcription velocity distribution will consequently also be bimodal. Such bimodal transcription dynamics is also clearly evident in individually measured trajectories (**Fig. 2D**), as previously reported (9). To characterize the heterogeneity in transcription velocity, we analyzed two groups of trajectories conditioned on their estimated pause probabilities (i.e. *P*_total_ < 0.08 and *P*_total_ > 0.08, respectively). While RNAps in the first group show an average velocity of ∼10 nt/s (**Fig. 2D**,**E**; purple), those in the second group advance at ∼5 nt/s (**Fig. 2D**,**E**; grey). Interestingly, we also detected a small fraction of RNAp (<5%) in which the velocity alternates between these two mean values (**Fig. 2D**,**F**), a phenomenon indicative of state-switching (34, 35). Together with the lack of clear correlation between *k*_el_ and *P*_total_ for individual trajectories (**Fig. S3A**), these results imply that variations in the frequency of short pauses, and not changes in *k*_el_, underlie the observed heterogeneity in transcription velocities.

We next asked how representative the combined DWT distribution is for the dynamic behavior of individual RNAps. We found that although *k*_el_ and *P*_total_ exhibit rather wide distributions, the combined DWT distribution that includes all trajectories (**Fig. S3D**) can be described impressively well by their average values. This establishes that, despite the inherent heterogeneity found in model parameters, they exhibit a self-averaging property in the sense that fitting the combined DWT distribution results in a population-averaged estimation of the parameters.

### All pauses compete with nucleotide addition

To validate the structure of our working model (**Fig. 2**), we first wished to establish whether all three identified pause states compete with the active catalytic pathway. To do so, we measured the response to [NTPs] ranging from 1 µM to 500 µM at a constant force of 7.5 pN AF. Overall, in agreement with previous results obtained from bulk and comparable single-molecule experiments, we found that a deficiency of NTPs reduced the average transcription velocity significantly (**Fig. 3A**), resulting in a proportional decrease in the processivity of single TECs (**Fig. 3B**) due to the statistically unchanged TEC lifetime (**Fig. S4A**) (9, 22, 31). The set of combined DWT distributions (**Fig. 3C**) present a more detailed view of the effect of NTP availability on the transcription dynamics. As the [NTPs] decreases, the elongation peak shifts to larger DWTs, indicating a slower *k*_el_. At the same time, the probability of all pauses gradually increases, suggesting that all three pauses are off-pathway states in (possibly indirect) competition with nucleotide addition. For the working model (**Fig. 3**, rightmost panel) to be true, the data set obtained at different [NTPs] should be fully captured by only allowing *k*_el_ and *P*_total_ to vary while keeping all other parameters fixed (**Table S2**). As a validation, we used this approach (**Methods M8**) to fit our model to the empirical DWT distributions measured at decreasing [NTPs] (1 µM - 500 µM). The resulting two-parameter fits (**Fig. 3C**, solid lines) fully capture the observed NTP-dependent variations in the combined DWT distributions.

We next investigated the NTP-concentration dependency of both *k*_el_ and *P*_total_. We here note that the mere existence of a competing pause branch affects the lifetime associated with the catalytic pathway - a fact largely ignored in previous studies. Taking this into account (**Methods M9**), the effective elongation rate *k*_el_ is given by

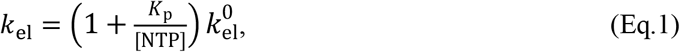

where the term in the parentheses represents the effect of the pause branch, with *K*_p_ given by **Eq. S34**, and 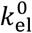 follows the Michaelis-Menten kinetics expected in the complete absence of a pause branch (14)

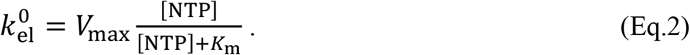

Equations **Eq.1** and **Eq.2** imply that *k*_el_ saturates at a finite value at very low [NTPs]. This can be understood by considering that 1/*k*_el_ represents the lifetime of the on-pathway elongation conditioned on *not* pausing: even if the nucleotide addition rate is very low, only elongation steps fast enough to occur prior to pausing will contribute to the lifetime. The value of *k*_el_ remains finite for the contributing subpopulation, while the size of the population shrinks as the pause probability approaches unity at the lowest [NTPs],

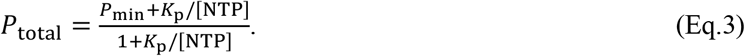

In the above, the parameters *V*_max_, *K*_m_, *K*_p_, and *P*_min_ depend only on the rates that constitute the active pathway and on the entry rate into the pause branch (**Fig. S4B**; **Methods M9**). Their precise values are expected to vary for different nucleotides (9, 36, 37). We found that the values of *k*_el_ and *P*_total_ estimated by the fits are well captured by **Eqs. 1** and **3** (**Fig. 3D**,**E**), leading to averaged (over 4 nt, based on the applied DWT-window) estimations of *V*_max_ = 21.8 ± 2.4 nt/s, *K*_m_ = 9.8 ± 2.5 *μ*M, *K*_p_ = 8.6 ± 2.8 *μ*M, and *P*_min_ = 0.07 ± 0.04. The model remains consistent with the empirical DWT distributions even if *K*_m_ is increased to 100 µM, compatible with *K*_m_ values previously measured for different individual nucleotides. Thus, our estimated value of *K*_m_ should be considered as a lower bound for the averaged Michaelis-Menten constant. In this case, deviations between the data and model predictions only occur in the left tail of the DWT distributions (**Fig. S4C**; i.e. at DWTs <0.2 s), which is most sensitive to (and can be biased by) experimental noise.

The fact that our estimations for *k*_el_ and *P*_total_ follow theoretical predictions (**Fig. 3D, E**) confirms that the model with three serially connected off-pathway pauses branching from the active catalytic state is compatible with our data, supporting the viewpoint that the elemental pause represents an off-pathway intermediate state that can isomerize into stabilized, long-lived pauses (2, 5).

### Recovery from deep backtracks is dominated by intrinsic cleavage

We next investigated the origin and interconnection of pause states. Previous single-molecule and bulk studies associated longer pause lifetimes with diffusive return from backtracking (11, 12, 19–22), which should be susceptible to force. We systematically measured the transcription dynamics of RNAp at applied forces ranging from 5 to 12.5 pN, in both the AF and OF directions (**Fig. S1A**) at 1 mM [NTPs]. As the magnitude of the OF is increased, we globally observe a substantial decrease in the average end-to-end velocity (**Fig. S5A**), indicating a change in either *k*_el_, *P*_total_, pause lifetimes, or a combination thereof. **Figure 4A** depicts examples of combined DWT distributions measured at 12.5 pN in AF and OF orientations (DWT distributions for all applied forces are shown in **Fig. S5B, C**). We found that switching the force direction from AF to OF produces a significant effect similar to that observed at low [NTPs]: enhanced pausing probability accompanied with a decrease of the effective elongation rate.

**Figure 4:**
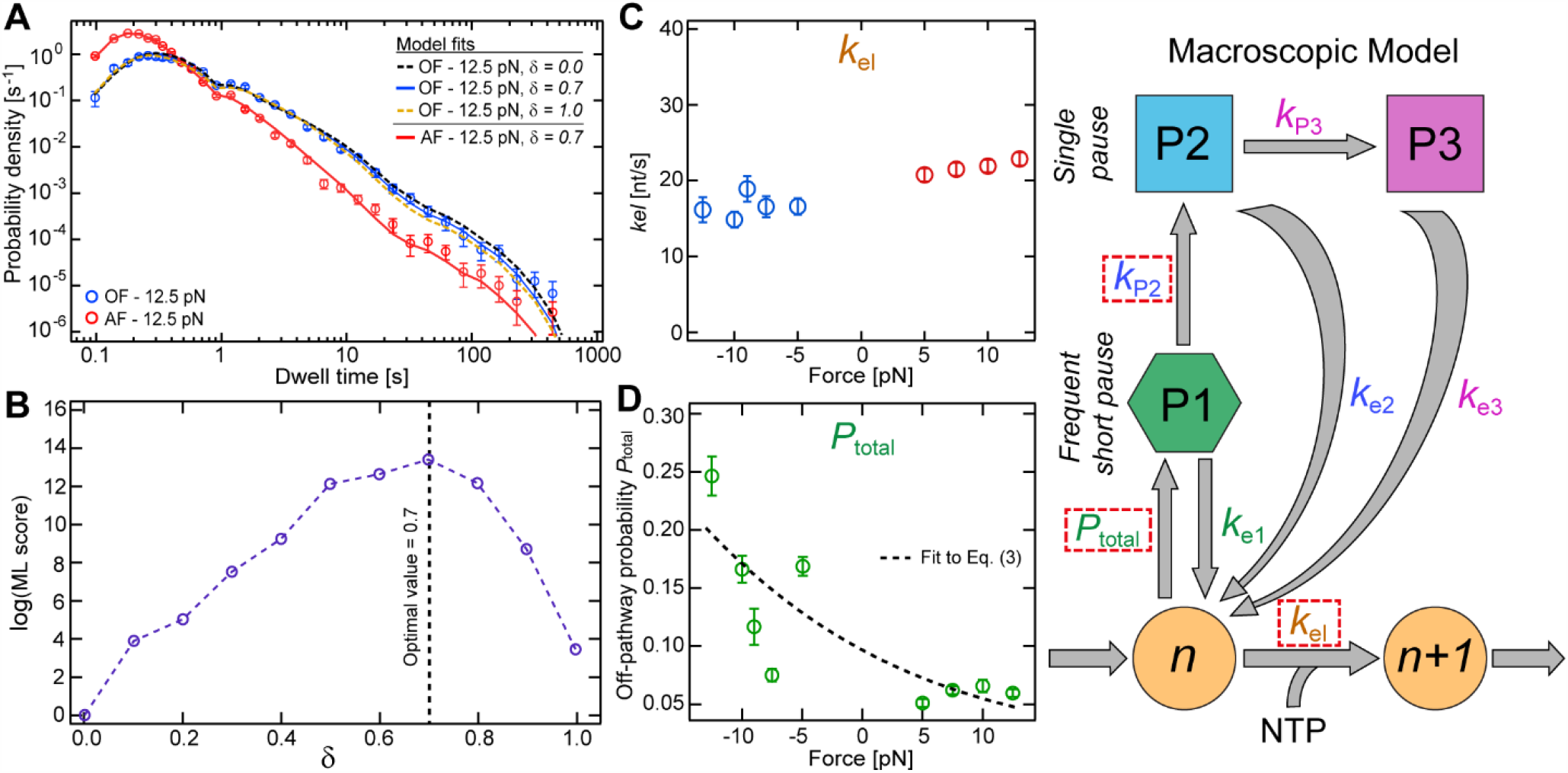
RNAp backtracking exhibits force-dependency in the entry rate, but not diffusion-governed recovery. (**A**) Combined DWT distributions (circles) measured at 12.5 pN AF and OF. The (dashed) lines represent the best fits of the model (rightmost panel) to each data set. Three fit parameters, *k*_el_, *P*_total_, and *k*_P2_, assumed force-dependent by the parameter *δ* (δ = 0: maximal force dependency, δ = 1: no force dependency; see Eq. 4), were used. (**B**) Fit quality as a function of *δ*, from fitting the OF 12.5 pN data set in (A). The maximum at 0.7 marks the optimal value for *δ* (**C** and **D**) The effective elongation rate *k*_el_ and pause probability *P*_total_, respectively, from the fits to data sets at different applied forces. The dashed black line in (D) depicts the theoretical prediction (Eq. 3) with *δ* = 0 7. **See also Figures S4, S5**, and **S6**.

To explain this force dependency, we first fitted our model (**Fig. 4**, rightmost panel) to the 12.5 pN OF data set by only allowing *k*_el_ and *P*_total_ to change while fixing all other parameters to the reference values (**Fig. 4A**, yellow dashed line). While the model fits the data at short timescales <10 s, it slightly deviates at longer timescales. This remaining discrepancy vanishes when we introduce force dependence in the entry rate of the P2 state as

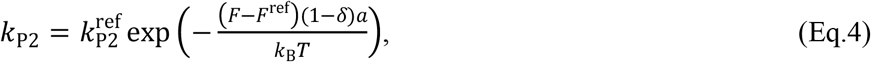

where *a* ≈ 0.37 nm is the average step size of RNAp, 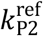 is the value corresponding to the reference force *F*^ref^ = 7.5 pN, *δ* identifies the position (in nucleotides) of the transition state between pre-translocated and backtracked states, and *F* is the applied external force (19, 38). By sweeping the range for *δ* between 0 and 1 (**Methods M8**), we found that the entry to P2 exhibits force-dependency with an optimal value of *δ*∼0.7 (**Fig. 4B**). Correspondingly, maximizing the force dependency of *k*_P2_ by setting *δ* = 0 results in an overestimation of the probability of the long-lived pauses P2 and P3 (**Fig. 4A**, black dashed line).

To validate proposed force-dependency for *k*_P2_, we performed two-parameter fits to the data sets obtained at different forces by using *k*_el_ and *P*_total_ as fit parameters, fixing *k*_P2_ at its expected value for each force assuming *δ* = 0.7 (**Eq. 3**; **Fig. S5D**) and fixing all other model parameters to their reference values (**Table S2**). The resulting fits to the data at 12.5 pN AF (**Fig. 4A**, red solid line), as well as to all other measured forces (**Fig. S5B**,**C**), demonstrate that our model remains valid at different forces.

We next explored the force dependency of the catalytic pathway. Each NMP addition cycle involves transition of RNAp from pre-to the post-translocated state, which is expected to be affected by an externally applied force (9). We found that *k*_el_, as estimated by our fits, remains fairly constant with force (**Fig. 4C**), implying that at saturating 1 mM [NTPs] RNAp downstream translocation does not become rate limiting across the studied force range, even at 12.5 pN OF. In contrast, the off-pathway pause probability *P*_total_ shows an increasing trend as the force is gradually swept from AF to OF (**Fig. 4D**). This important observation indicates that pausing is in competition with TEC translocation, justifying our assumption that the entire pause branch originates from the pre-translocated state.

Using **Eq. S36** to describe the force dependency of *P*_min_ and *K*_p_ in **Eq. 3** (**Methods M9**), with the reference values 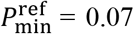 and 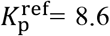 corresponding to the force *F*^ref^ = 7.5 pN (**Fig. 3E**), we fitted **Eq. 3** (**Fig. 4D**, dashed line) to the estimated values of pause probability (**Fig. 4D**, circles). The best fit corresponds to Δ ≈ 0.5 ± 0.16, which identifies the position of the transition state against RNAp transition from the pre-to post-translocated state (**Eq. S35**). We note that around 5 pN OF a non-monotonic variation of the pause probability provides some discrepancy with the model prediction. Possibly, this results from an inhomogeneity in the magnitude of force applied to the different magnetic beads (39).

The entry rate *k*_P2_ into the first longer-lived pause state P2 increases with OF (**Fig. S5D**), prompting us to propose that P2, and in consequence, P3 are backtracked pauses. RNAp can stochastically switch to a catalytically-inactive backtracking mode, where it can diffuse back and forth along the template DNA (40, 41). In a backtracked TEC, the nascent RNA is threaded through the active site, blocking nucleotide addition. The RNAp can recover from backtracking either by diffusing forward, placing the 3’end of the transcript in the active site, or by cleaving the backtracked RNA, thereby freeing the active site (16). Previous single-molecule studies have suggested that diffusion provides the dominant recovery mechanism from long-lived pauses (11, 12, 19–22).

We thus investigated whether the force dependency reflected in our data can be explained by diffusive backtrack recovery. Evident from our analysis, and in agreement with other reports (20, 22), the P2 and P3 lifetimes are insensitive to external forces. This effect has been previously rationalized in terms of a force-independent *t*^−3/2^ power-law region in the DWT distribution, a signature attributed to diffusive backtrack recovery (19). This power-law behavior, however, will eventually transform into an exponential cut-off at large external forces or long observation times. Such an exponential cut-off is clearly apparent in our data at extended DWTs (**Figure 4A**; **Fig. S5B, C**), which motivated us to assess (**Methods M10**) the compatibility of the diffusive backtracking model over the full temporal range of our data. The result of this analysis is depicted in **Figure 5**. We found that the diffusive backtracking model captures the combined DWT distribution of the reference data well, provided that the maximum backtrack depth (BT) exceeds 5 nt (**Fig. 5A)**, which is compatible with previous studies that assumed unrestricted diffusive backtracking to describe long-lived pauses. However, the agreement between the data and the diffusive backtracking model fails to persist at high OFs, and none of the simulated DWT distributions, including the one with the highest likelihood score (**Fig. 5B**, red line), satisfactorily captures the data. Moreover, the simulated trajectories corresponding to the best fit typically show a significantly shorter processivity than the measured data, an inherent property of the diffusive backtracking model in which at sufficiently high OFs eventually all RNAps will be trapped in a deep backtrack (**Fig. 5C**) (19). We found that this occurs in the diffusive backtracking model provided that the maximum BT exceeds 5 nt, leading to simulated trajectories which show, on average, a processivity at least one order of magnitude lower than the mean value deduced from our measured RNAp trajectories within >2,000 s of observation time (**Fig. 5D**). Thus, while the diffusive backtracking model can be fitted to any individual DWT distribution at a constant force, it does not provide a consistent description of our data over the entire force range and significantly underestimates RNAp processivity. Importantly, this strongly indicates that diffusive RNAp dynamics alone cannot account for the observed long-lived pauses in our empirical DWT distributions.

**Figure 5:**
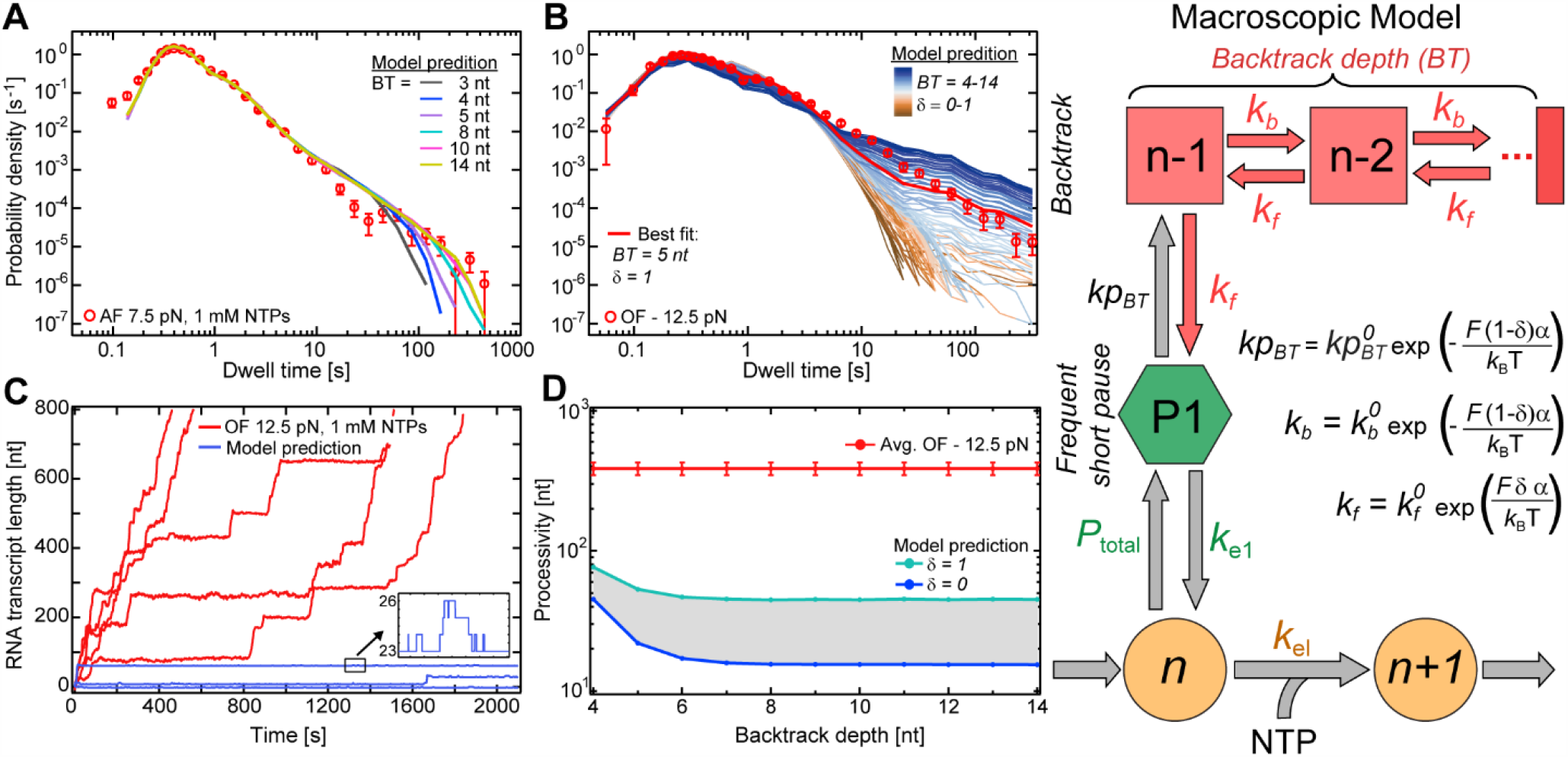
RNAp backtrack-recovery is non-diffusive. (**A**) DWT distribution (red circles) of reference data measured at 7.5 pN AF, 1 mM NTPs. The lines correspond to the best fits achieved using the diffusive backtracking model (top right panel) to fit the data, where the maximum backtrack depth (BT) was restricted to 3 - 14 nt. **(B**) DWT distribution (red circles) data measured at 12.5 pN OF, 1 mM NTPs. The lines represent the ML fits of the diffusive backtracking model (rightmost panel) to the data, with different BT values, with the Pause 2 entry rates (*k*_P2_) described by the parameter *δ*, and all other kinetic parameters as fixed to the values obtained from the fit to the data measured at AF 7.5 pN (Figure 2A). The red line represents the best fit, with *δ* = 1 and a maximum BT of 5 nt. (**C**) Measured (red lines) and simulated (blue lines) trajectories of RNAp under 12.5 pN OF over time. The used model parameters correspond to the best fit to the OF data (red line in B). (**D**) Average processivity for measured data (red; 12.5 pN OF, 1 mM NTPs) compared to the model prediction for different values of maximum BT and *δ*. The lines mark the minimum (*δ* = 0; blue) and maximum (*δ* = 1; cyan) average processivity at each BT, as predicted by the model. Any other choices for *δ* lead to a processivity in between the two limits (gray area). **See also Figure S6**.

The intrinsic RNA cleavage provides an alternative pathway for recovery from backtracking by allowing RNAp to restart RNA chain extension upstream from the position where backtracking was initiated (**Fig. S6A**). Although an external force can affect the backtrack depth reached before cleavage, if the time for a TEC to return to its original position were small compared to a typical cleavage time, such a recovery mechanism would be compatible with the observed lack of sensitivity of the P2 and P3 lifetimes to external force. Even at the lowest estimated effective elongation rate (17 nt/s at 12.5 pN OF), this condition approximately holds for backtrack depths up to about 20 nt, as it takes <1 s on average to translocate back to the original position (**Fig. S6B**). Therefore, we take both the force independence and the incompatibility of the diffusive backtrack model with our data as indications that intrinsic cleavage forms the main recovery pathway from deep backtracks.

### Deep backtracked pause states are affected differently by Gre cleavage factors

We probed the contribution of RNA cleavage to backtrack recovery dynamics in the single-molecule context using transcription elongation factors GreA and GreB, which augment the intrinsic RNA cleavage rate in backtracked TECs (42, 43).

To increase the measurable effect of Gre factors, we applied an OF that favors the long-lived pauses (**Figs. 4** and **S5**). **Figure 6A** shows the effect of 2 µM GreA and 2 µM GreB have on the DWT distribution measured at 9 pN OF, 1 mM [NTPs]. It is evident that solely pauses exceeding 4 s, corresponding to the P2 and P3 states (**Fig. 2A, B**; **Table S2**), are affected by both Gre factors, and that the effect is considerably more pronounced for GreB. We fitted our model (**Fig. 6**) to the Gre factor data sets (**Methods M8**) by only allowing *k*_e2_, *k*_P3_, and *k*_e3_ to vary; all other model parameters were fixed according to the best fit to the corresponding control data acquired in the absence of Gre factors. We found that the three-parameter fits were able to fully capture the effects of both Gre factors (**Fig. 6A**); the resulting fit values are shown in **Figure 6B, 6C**, and **6D**, respectively. Compared to the control data, the effect of GreA was statistically insignificant, while GreB significantly increased *k*_e2_ while leaving *k*_P3_ and *k*_e3_ unchanged within errors.

**Figure 6:**
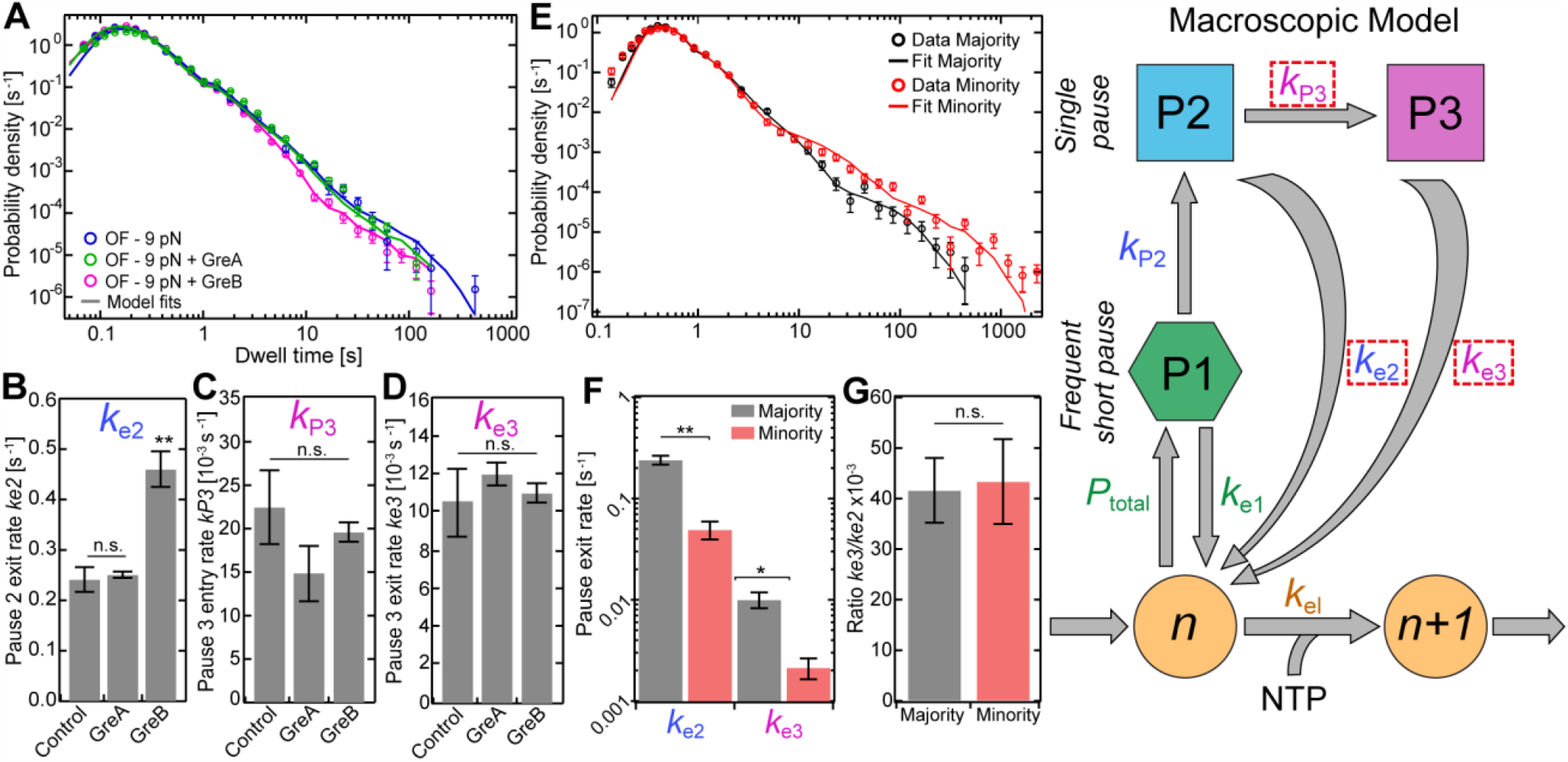
GreA/B-mediated cleavage and heterogeneity in intrinsic cleavage. (**A**) Superimposed combined DWT distributions (circles) of trajectories measured at OF 9 pN in presence of GreA (green) or GreB (magenta), and absence of either (blue). The lines in (A) represent the fits of our model (rightmost panel) to each data set using *k*_e2_, *k*_P3_ and *k*_e3_ as the only free fit parameters (dashed rectangles). (**B**) *k*_e2_, (**C**) *k*_P3_, and (**D**) *k*_e3_ estimated from the fits in (A). (**E**) Superimposed DWT distributions (circles) and fits (lines) for the minority and dominant sub-populations, extracted from the reference data set and separated by Gaussian mixture clustering (Methods M4) with respect to their maximum pause durations. (**F**) *k*_e2_ and *k*_e3_ for the majority (grey) and minority (red) sub-populations, estimated from the fits in (E). (**G**) The ratio between *k*_e2_ and *k*_e3_ for the dominant (grey) and minority (red) sub-populations. Statistical analyses consisted of unpaired, two-tailed t-tests (significance level p: ** ≤ 0.01, * = ≤ 0.05; n.s. = non-significant). **See also Figures S4** and **S6**.

Since GreB is known to facilitate recovery from deep backtracks (42, 43), the observed acceleration of the exit rate *k*_e2_ indicates that the P2 pause originates from backtracked TECs. The concomitant reduction in the occurrence of the P3 pause state, reflected by a decrease in the population of pauses longer than 10 s in **Figure 6A**, supports our assumption that the entry to P3 competes with recovery from P2, implying that P3 also originates from a backtrack-induced pause. Backtrack depth analysis also reveals that P2 and P3 are associated with backtracks with similar depths >4 nt (**Fig. S6A**,**B**). In contrast to P2, the lifetime of P3 (**Fig. 6D**) is largely unaffected by GreB. This suggests that to enter P3, P2-paused RNAp may undergo conformational changes that render it resistant to GreB-assisted cleavage, which in turn leads to a 20-fold slower recovery rate as compared to the recovery from P2.

Our observation that GreA has a negligible effect on short pauses is in agreement with previous single-molecule studies (44, 45) and could originate from the fact that it mainly affects backtracks of ≤3 nucleotides (43). This result indicates that shallow backtracks are short-lived and either recover quickly (≤1 s) *via* rapid intrinsic cleavage or diffusion, or lead to extended backtracking. This scenario could explain why short backtracks remain undetected in our analysis, as their contribution to the DWT distribution might be overshadowed by the elemental pause (5). Furthermore, if short backtracks are predominantly recovered by diffusion (which our data do not report on), GreA-mediated cleavage might not be able to substantially suppress the occurrence of deeper backtracks.

### A minority population reveals the nature of long-lived backtrack-associated pauses

As described above, variations in the duration of long pauses between individual RNAp trajectories (**Fig. 1D**) allowed us to distinguish (**Methods M4**) a dominant sub-population (∼90%), to which we fitted our working model, and a minority sub-population (∼10%). The latter population contained trajectories with pauses exceeding thousands of seconds and has thus far been excluded from our analyses. Here, we ask whether the RNAp dynamics in this minority sub-population can be understood in the context of our model.

We constructed a combined DWT distribution using only the minority sub-population from the reference data set (**Fig. 6E**, red circles), and performed a maximum likelihood fit (**Fig. 6E**, red line) to obtain a new set of model parameters. We found that only the exit rates from P2 and P3 vary significantly between the minority and majority sub-populations (**Fig. 6F**), while all other parameters remain statistically unchanged (**Fig. S6C**). We note that the P2 lifetime is shorter than the 450 s cutoff used in the selection process, and thus is not affected by the RNAp trajectory selection based on the duration of the longest pause. The fact that not only long P3 pauses but also shorter P2 pauses are different between the two sub-populations therefore confirms that differences in their exit rates reflect an actual change in RNAp dynamics rather than being artifacts originating from the imposed cutoff in our selection process. Strikingly, the ratio between these exit rates is comparable in both sub-populations (**Fig. 6G**), suggesting that rate-limiting sub-processes that form the P2 recovery pathway also dominate the P3 recovery, consistent with TEC escaping from both pause states by cleavage. While P2 and P3 do not appear to differ in backtracking depth (**Fig. S6B**), recovery from P3 occurs ∼20 times more slowly and cannot be accelerated by GreB. These observations are consistent with P3 resulting from an infrequently occurring, long-lived conformational change in backtracked TEC that must be reverted prior to RNA cleavage.

## DISCUSSION

Based on global *in silico* modelling of high-throughput data sets collected over many experimental conditions, we have provided a complete quantitative characterization of the full temporal spectra of RNAp pausing dynamics at the single-trajectory level. We have established the hierarchical relations between transcriptional pauses and their origin, which confirm and unify many previous observations made in isolation. Through this study, we have provided distinct evidence that the observed heterogeneity in average transcription velocity and the previously reported dynamic state switching are related, both originating from variations in the frequency of short transcriptional pauses. Furthermore, we have identified a cleavage-deficient subpopulation of RNAps which exhibit long-lived backtrack pauses. The characterization of this subpopulation, together with probing the effect of Gre factors on transcriptional pausing dynamics, has allowed us to identify two distinct backtrack pause states which differ in the intrinsic cleavage rate of RNAps. This discovery has allowed us to further elucidate the mechanism of recovery from deep backtracks and rationalize the contradicting viewpoints presented in the literature regarding the RNAp backtracking dynamics.

### A unifying mechanistic model of intrinsic transcription dynamics

Based on our findings, we propose a full kinetic model (**Fig. 7**) in which frequent short-lived pauses, consisting of the elemental pause (EP) and shallow backtracks (BT) of lifetimes <1 s, lead to two long-lived pauses (P2 and P3) accompanied by deeper backtracks >4 nt.

**Figure 7:**
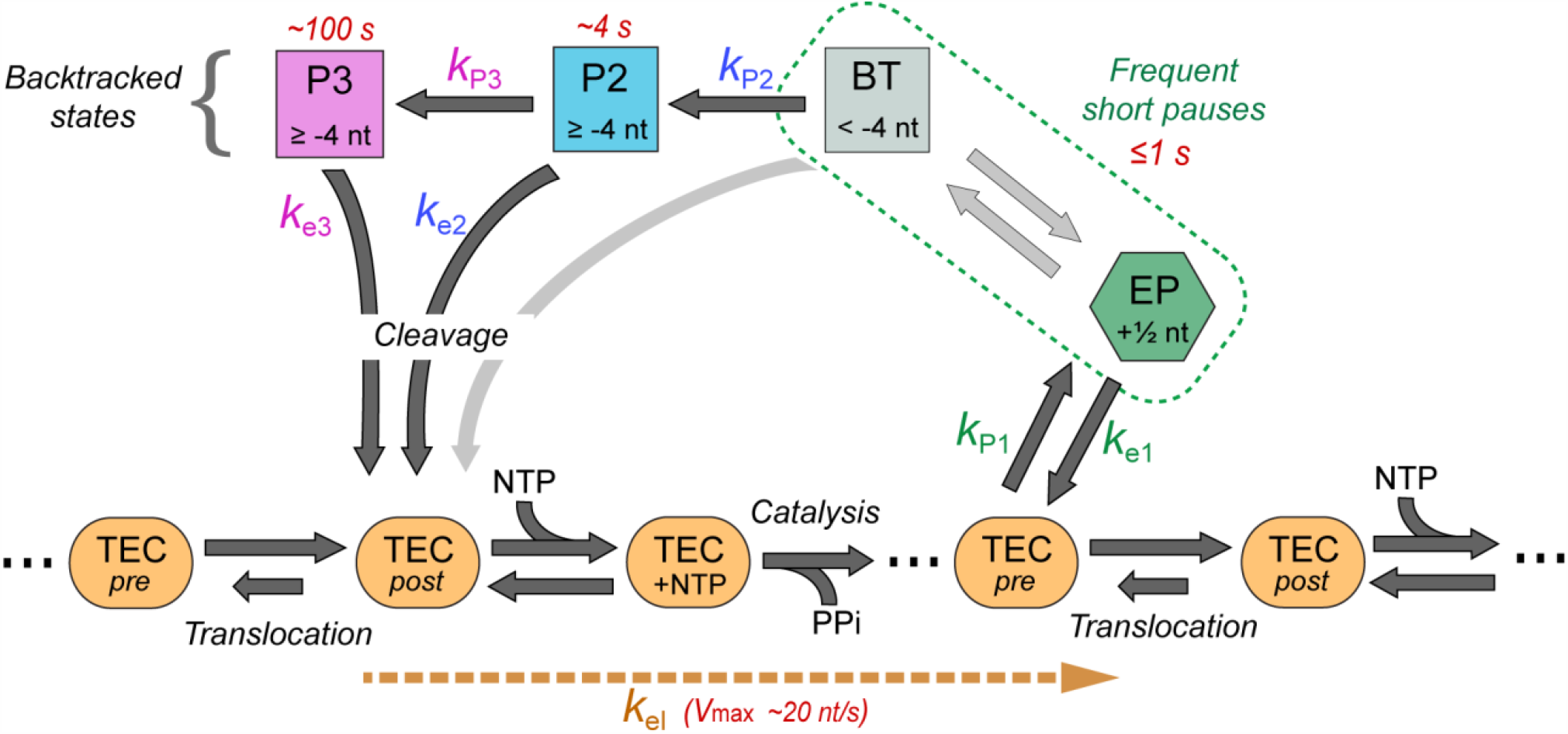
Unified mechanistic model of transcription. The active pathway includes the pre-translocated (“*pre*”), post-translocated (“*post*”), NTP-bound (“*+*NTP”), and catalysis (with PP_i_ release) states of the TEC. The sub-processes involved in RNA elongation, translocation, NMP-addition, and catalysis, collectively contribute to the effective elongation rate *k*_el_. The pause pathway includes the - half-translocated - elemental pause (EP) (5), the -undetected-fast pause (BT) associated with backtracks <-4 nt and two long-lived pauses (P2 and P3) associated with deep backtracks with cleavage-mediated recovery. For each pause state, the assumed position of the RNAp’s active site with respect to the pre-translocated state is identified. The estimated maximum velocity (*V*_max_; Eq. 1) and lifetimes for each pause state, for the dominant sub-population, are denoted in red.

Our results are largely consistent with previous models of RNAp that have assumed serially connected pause states (5, 18, 38, 46). The shortest pause state, with a lifetime ≤1 s, resembles the elemental pause identified in previous studies that consists of a half-translocated state (5, 14). We showed that the two long-lived pauses are promoted by OF and thus represent RNAp backtracking, but that their lifetimes are not affected by force. In previous studies, this lack of force dependency has been commonly attributed to a force-independent power-law tail of the DWT distribution that originates from diffusive backtracking (9, 11, 12, 19–22, 31, 44). Using a longer observation time, we were able to discriminate power-law behavior from exponentially distributed states. We found that the diffusive backtracking model does not provide a consistent description for our data across the entire force range. Instead, our findings imply that the RNAp recovers from the two long-lived pauses, P2 and P3, by RNA cleavage with rates comparable to previous biochemical studies (5, 15, 16, 33, 42). Based on the observation that these pause states differ in their susceptibility to intrinsic and Gre-assisted RNA cleavage, we established that both P2 and P3 correspond to backtracks deeper than approximately four nucleotides, and propose that they are separated by a conformational change in RNAp as further discussed below. The fact that backtracks shorter than ∼4 nt remain undetected in our analysis implies that their recovery, whether through diffusion or cleavage, is faster than or comparable to the elemental pause.

The proposed kinetic model with serially connected pause states, and RNA cleavage as the predominant recovery mechanism from long-lived deep backtrack states, largely resembles the observations made for Pol II (11, 18, 47, 48). After entering the ubiquitous catalytically inactive elemental pause state, RNAp, as well as Pol II can both transition into backtracked states where the intrinsic cleavage acts as a potential recovery mechanism in kinetic competition with the Brownian diffusion. While both polymerases return from shallow backtracks mainly by diffusion, the intrinsic cleavage becomes dominant as the backtracking depth increases. Deep backtracks cause a temporary transcription arrest, from which recovery is only possible by intrinsic cleavage of the transcript or by cleavage in conjunction with elongation factors. The observation that the RNAp P2 and P3 pause states have vastly different lifetimes is similar to the proposed model for Pol II with cleavage rate spanning a similar time range depending on the backtrack depth.

Our proposed consensus mechanistic model that unifies our and previous key findings will serve as a baseline for future studies on bacterial RNAP, allowing for a quantitative characterization of effects of pause-inducing sequences, transcription factors, RNAp-targeting antimicrobials, among others, on transcription dynamics. By applying the statistical analysis presented in this work to such studies, and comparing the outcomes to this baseline, it would be possible to identify the RNAp kinetic state(s) the above-mentioned elements act on, as well as any newly induced kinetic states.

### The nature of long-lived backtrack-stabilized kinetic states

Our analysis revealed that *E. coli* RNAp can fall into long-lived, cleavage-resistant pause states while actively transcribing a template that is free from known pause-inducing sequences (10, 12), as similarly observed in previous studies. Although TECs with similar properties have been documented in biochemical studies (16, 49, 50), these complexes were scaffolds assembled and equilibrated for an extended time in an arrested state. Remarkably, the rate of RNA hydrolysis in *E. coli* TECs can vary over 60,000-fold depending on the sequence context and the presence of Gre factors (16), with TECs assembled on scaffolds containing an elemental pause sequence or a deeply (∼9 bp) backtracked RNA being particularly resistant to intrinsic and Gre-stimulated cleavage (16, 49). The arrest of RNAp in a cleavage-resistant state has also been observed in single-molecule investigations, where stalling of RNAp through applied force were found to be unable to resume transcription, even in presence of GreB (31, 51), suggesting that conformational changes in the active site cause the mechanistically arrested state. Conformational transitions of several mobile RNAp domains, most notably the β’ subunit catalytic trigger loop (TL) and a large *E. coli*-specific SI3 insertion in the middle of TL, have been proposed to explain these differences (16, 49, 50).

Recent cryo-EM structures of *E. coli* TECs stabilized in a paused state upon formation of an RNA hairpin (52, 53) or by an engineered backtrack (4) confirmed these predictions. In inactive TECs, *E. coli* RNAp undergoes a rotational motion termed *swiveling*, which repositions the TL and SI3 modules, preventing the TL folding and thus inhibiting catalysis. A similar ratcheting motion accompanies the formation of a backtracked *Thermus thermophilus* TEC (17). By contrast, catalytically-competent TECs, including a GreB-bound backtracked TEC poised for RNA cleavage (4), are not swiveled.

Swiveling is thought to accompany the formation of long-lived paused states at regulatory pause sites, such as hairpin-stabilized *his*P (2). Interestingly, our analysis revealed two distinct long-lived paused states, both of which are eventually able to resume transcription but differ significantly in their sensitivity to RNA cleavage. These results suggest that RNAp may occasionally swivel even on non-pause sequences, assuming an inactive conformation. Since we and others (9, 16, 20, 54, 55) failed to observe significant differences in the backtrack depth between GreB-sensitive (P2) and -resistant (P3) states, we hypothesize that, independent of the backtrack depth, P2 TECs can either stochastically revert to the catalytic pathway through RNA cleavage or undergo additional conformational changes into the inert P3 state. Our results indicate that majority of TECs that remain in the P2 state longer than 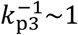 minute will be found in such inert P3 state. This estimation is largely supported by studies that ‘walked’ TECs into backtracked states of various depths and stalled the RNAp for several minutes, reporting cleavage times comparable to our estimation of the P3 lifetime (∼100 s, up to ∼200 s).

While the triggers and structural changes that accompanies P3 formation are not known, repositioning of the β’ rim helices or SI which directly contact GreB (4) or slight changes in the path of backtracked RNA (55) would strongly affect endonucleolytic cleavage. The free RNA originating from deep backtracks could also act as a potential trigger by forming secondary structures, constraining diffusional return and thereby inducing conformational changes into an inert state. Recovery from the inert P3 state would then require reversal of such additional changes and counter-swiveling into an active conformation. Future structural studies, focusing on TEC stalling and factor-assisted cleavage, may provide further insights into the properties of different backtrack states.

### The origin of heterogeneity in transcription velocity and pausing dynamics

The majority of previous studies aggregated all measured RNAp transcription trajectories based on the assumption that all RNAp have similar dynamics; yet evidence in support of dynamic state switching, which involves alternation between different transcription velocities of a single RNAp, has also emerged (23, 34). However, most studies failed to detect state-switching, concluding that individual RNAps transcribe at a constant velocity, with apparent differences possibly due to arbitrary structural and chemical variations between individual RNAps that arise during their expression and holocomplex assembly (32, 56).

By capturing the full extent of temporal transcription dynamics with our large ensemble of RNAp trajectories, we observed considerable heterogeneity in elongation rates and pause frequencies for individual RNAp. Particularly, while the majority of RNAps transcribed at similar average transcription velocities (∼10 nt/s), a notable fraction of RNAp (∼10%) transcribed the template more slowly (∼5 nt/s), and a small fraction (∼5%) of RNAp trajectories showed switching between these two velocities. Strikingly, these velocities are quantitatively compatible with the previously described state switching (34). The characterization of the transcription dynamics of identified RNAp trajectories with different average velocities indicated that state switching originates from alterations in the frequency of short pauses, since the effective elongation rate remained unaffected. We speculate that the observed variation in pause frequency may reflect conformational changes in one or more mobile modules of RNAp. Most notably, dynamics of two β’ elements, the TL and the bridge helix (BH), are central to catalysis and translocation. Upon cognate NTP binding, TL refolds into trigger helices (TH) and forms a three-helix bundle with the BH to position the 3’ OH and the NTP substrate facilitating catalysis (57); the TH unfolds into the TL prior to the next nucleotide addition cycle. The TH contains two flexible hinges (58, 59). Altered motions of TH hinges may reduce RNAp velocity to the ∼5 nt/s observed here; e.g., since locking the folded TH by crosslinking has shown to reduce the elongation rate (60). Since the described transcription heterogeneity persists throughout our experiments and is reversible in some cases, we speculate that cis-trans isomerization of the universally conserved Pro residues in TH hinges may underlie this phenomenon. Alternatively, locally misfolded states of BH, TL, or other mobile domains could hamper their concerted transitions and thus prevent rapid elongation.

We have further identified an additional source of heterogeneity linked to significant differences in long-lived pause lifetimes associated with backtracked TEC. A minority sub-population of RNAp (∼10%) showed notably slower backtrack-recovery rates from both the P2 and the P3 states, indicating a slower intrinsic cleavage rate. We note that the longest pause durations in this sub-population exceed thousands of seconds, and have been previously categorized as arrested states unable to recover from backtrack (16, 20, 21, 31, 34, 61). Since many structural elements involved in RNA synthesis also participate in the RNA cleavage, we assume that the suggested possible chemical modification of different structure motifs during RNAp synthesis may also be responsible for the observed RNA cleavage-deficient sub-population. Elucidating the microscopic origin and cause of the observed different dynamic RNAp behavior remains an important subject for future biochemical and structural studies.

## General

We thank Theo van Laar for DNA construct synthesis, Nina Turk for assistance in single-molecule measurements, Mariana Köber for backtrack depth analysis, and Georgiy Belogurov for discussions.

## Funding

Funding to N.H.D. was provided by the Netherlands Organization for Scientific Research (NWO) via a TOP-GO program grant and a FOM Vrij Programma grant, and by the European Union via an ERC Consolidator Grant (DynGenome, no 312221). Funding to I.A. was provided by the National Institutes of Health grant GM67153. B.E.M. forms part of the research program ‘‘Crowd management: the physics of genome processing in complex environments”, supported by the Netherlands Organisation for Scientific Research (NWO).

## Author Contributions

R.J., B.E.M., I.A., M.D., and N.H.D. designed the experiments. R.J. performed the single-molecule experiments and processed data. B.E.M and M.D. designed the analysis pipeline, and B.E.M. performed the dwell-time analysis and *in silico* modeling. All authors contributed to the written manuscript.

## Competing interests

The authors declare no competing interests.

## SUPPLEMENTARY INFORMATION

Supplementary information for this article includes 6 figures and 3 tables, and is available at http://

## MATERIALS AND METHODS

### Purification of *E. coli* RNA polymerase holoenzyme and Gre factors A and B

Wild-type *E. coli* RNA polymerase holoenzyme with pre-bound transcription factor σ^70^ was purified as previously described (62). The enzyme contains a biotin-modification at the ß’-subunit that serves as an anchor to attach streptavidin-coated magnetic beads (9). GreA and GreB factors were individually obtained and purified following a previously established protocol (63). The activity of all purified proteins was confirmed using standard bulk transcription assays.

### DNA constructs for single-molecule experiments

To create a digoxigenin (DIG)-enriched handle, a 643 bp fragment from pBluescript Sk+ (Stratagene, Agilent Technologies Inc., USA) was amplified by PCR in the presence of Digoxigenin-11-dUTP (Roche Diagnostics, Switzerland) using primers 1 and 2 (Table S3). Oligonucleotides (Table S3) were obtained from Ella Biotech GmbH, Germany.

For the assisting force (AF) configuration, the digoxigenin-enriched DIG handle was ligated to a 4015 bp spacer consisting of lambda phage sequence from the plasmid pblue1,2,4 + pSFv1A using primers 3 and 4 (Table S3) followed by the T7A1 promotor in front of the RpoB coding sequence and the T7 terminator derived by PCR using plasmid pIA146 and primers 5 and 6 (Table S3). This resulted in a linear dsDNA construct of 9.2 kb.

For the opposing force (OF) configuration, the T7 terminator site was removed from plasmid pIA146 by digesting the plasmid with *Hin*dIII and *Sph*I (New England Biolabs, UK). Blunt ends were created using the Klenow fragment of DNA polymerase I (New England Biolabs, UK), and these blunt ends were ligated together with T4 DNA ligase (New England Biolabs, UK), resulting in plasmid pIA146Δterminator. DIG handles were ligated to a 1268 bp PCR fragment from plasmid pIA146Δterminator using primers 7 and 8 (Table S3) and a 5543 bp PCR fragment from plasmid pIA146 containing the T7A1 promotor and the *E. coli* RpoB coding sequence using primers 9 and 10 (Table S3). Prior to ligations, all amplicons were digested with the non-palindromic restriction enzyme *Bsa*I-HF (New England Biolabs, UK). The ligation of the fragments was carried out using T4 DNA ligase (New England Biolabs, UK). This resulted in a linear dsDNA construct of 7.5 kb.

### M1: Single-molecule RNAp transcription assay

The flow cell preparation used in this study has been described in detail elsewhere (26). In short, polystyrene reference beads (Polysciences Europe) of 1.5 µm in diameter were diluted 1:1500 in PBS buffer (pH 7.4; Sigma Aldrich) and then adhered to the nitrocellulose-coated (Invitrogen) surface of the flow cell. Further, digoxigenin antibody Fab fragments (Roche Diagnostics) at a concentration of 0.5 mg/ml was incubated for ∼1 hour within the flow cell, following incubation for ∼2 hours of 10 mg/ml BSA (New England Biolabs) diluted in buffer A containing 20 mM Tris, 100 mM KCl, 10 mM MgCl_2_, 0.05 % (v/v) Tween20 (Sigma Aldrich) and 40 µg/ml BSA (New England Biolabs), adjusted to pH 7.9.

The preparation of the RNAp ternary complex was performed as described previously (9, 26). Briefly, RNAp holoenzyme was stalled on the DNA constructs at position A29 after the T7A1 promoter sequence. To do so, 30 nM of RNAp holoenzyme (with σ^70^) was added to 3 nM linear DNA template in buffer A and incubated 10 min at 37°C. Afterwards, 50 µM ATP, CTP, GTP (GE Healthcare Europe), and 100 µM ApU (IBA Lifesciences GmbH) were added to the solution and incubated for additional 10 min at 30°C. The ternary complex solution was diluted to a final concentration of 250 pM of the RNAp:DNA complex. The complex was flushed into the flow cell and incubated for 30 min at room temperature. The subsequent addition of 100 µL streptavidin-coated superparamagnetic beads (diluted 1:400 in PBS buffer; MyOne #65601 Dynabeads, Invitrogen/Life Technologies) with a diameter of 1 µm resulted in the attachment of the beads to biotinylated RNAp stalled on the DNA.

Transcription was re-initiated by adding ATP, CTP, GTP, and UTP (GE Healthcare Europe) to buffer A at equimolar concentration of 1 mM to the stalled RNAp ternary complexes and immediately starting the single-molecule measurements. The experiments were conducted for 2.5 hours at constant pulling forces with a camera acquisition rate of 25 Hz. Instrumental drift was excluded by the use of surface-attached reference beads.

Transcription traces were processed using custom-written Igor v6.37 and MatLab R2016b-based custom-written scripts. The absolute z-position of the RNAp during the transcription process was converted to transcribed RNA product as a function of time, using the end-to-end length determined by the extensible worm-like chain model with an experimentally determined stretch modulus of 800 pN and a persistence length of 56 nm (64). To reduce the effect of Brownian noise in the applied DWT analysis and MLE fitting, all elongation traces were filtered prior to 1 Hz using a sliding mean average filter.

### M2: Magnetic tweezers instrumentation

The magnetic tweezers implementation used in this study has been described previously (26). Briefly, light transmitted through the sample was collected by a 50x oil-immersion objective (CFI Plan 50XH, Achromat; 50x; NA = 0.9, Nikon) and projected onto a 12-megapixel CMOS camera (#FA-80-12M1H, Falcon2; Teledyne Dalsa) with a sampling frequency of 25 Hz. The applied magnetic field was generated by a pair of vertically aligned permanent neodymium-iron-boron magnets (Webcraft GmbH, Germany) separated by a distance of 1 mm, suspended on a motorized stage (#M-126.PD2, Physik Instrumente) above the flow cell. Additionally, the magnet pair can be rotated around the illumination axis by an applied DC servo step motor (C-150.PD; Physik Instrumente). Image processing of the collected light allowed us to track the real-time position of both surface attached reference beads and superparamagnetic beads coupled to RNAp in three dimensions over time. The bead *x, y, z* position tracking was achieved using a cross-correlation algorithm realized with custom-written software in LabView (2011, National Instruments Corporation) (25). The software determined the bead positions with spectral corrections to correct for camera blur and aliasing.

### M3: Statistical DWT analysis of RNAp transcription trajectories

The transcription dynamics of *E. coli* RNAp were quantitatively assessed using unbiased dwell time analysis (24, 26). In this approach, the times needed for RNAp to transcribe through consecutive dwell time windows of a chosen size - defined as *dwell times* (DWT) - were calculated for all RNAp trajectories and used to construct a DWT probability distribution function. Since the validation of different dwell-time windows does not affect the analysis (Methods M6, Fig. S2), we chose a DWT-window of 4 nt and data filtering to 1 Hz filtering for analysis of all data sets within this work. The expected error (standard deviation)in the constructed distribution was estimated by bootstrapping the data 100 - 1,000 times (24, 26, 29).

### M4: Selection of RNAp transcription trajectories based on Gaussian mixture model

To select a homogeneous ensemble, a preselection protocol was applied to each data set, where a Gaussian mixture model were used to classify the transcription trajectories into three clusters, based on the value of the largest pause (or DWT) that was measured for each trajectory (65). The trajectories associated to the cluster with the largest mean were excluded.

### M5: Decomposition of the theoretical dwell-time distribution into exponential processes

In this section an analytical expression is derived which is used to fit our proposed model to the reference data set. To achieve this, certain assumptions and approximations have to be made as described below. We validate these assumptions in section M7 using computer simulation, proving that the analytical expression indeed captures the data well.

Assuming a DWT-window of 1 nt, the DWT distribution as predicted by the model (Figure 2, rightmost panel), can be decomposed into four separate terms,

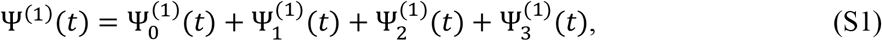

where 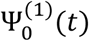 corresponds to RNAp taking a step without entering the pause pathway, 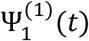 (*t*) includes entering P1 at least once without entering P2 or P3, 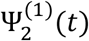 includes transitions into P2 without entering P3 and 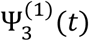 captures transitions into P3. Defining the exponential distributions,

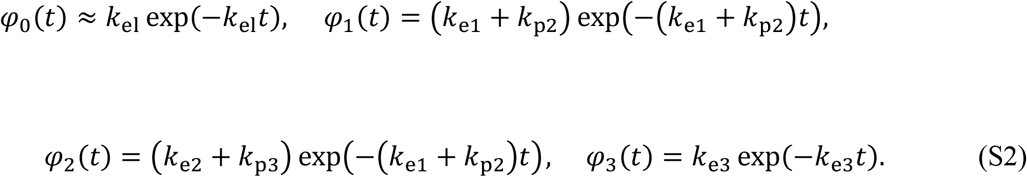

which describe the distribution of the lifetime of the various states in the model, as well as the splitting probabilities,

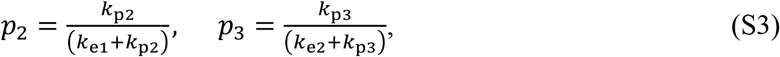

an analytical expression can be found for each component of **Eq**. (**S1**) in the Laplace space. Denoting the Laplace transformation of a function *f*(*t*) as 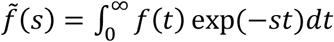, we have

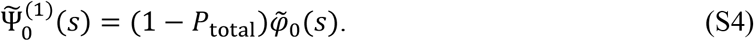

Furthermore, defining

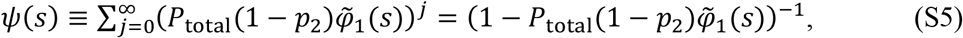

and assuming that the rate into the pause branch is much faster than the pause exit rates, one can write

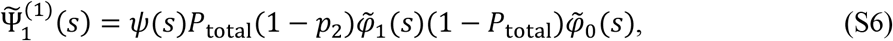

where the *j*_th_ term in the sum in **Eq**. (**S5**) accounts for *j* successive back-and-forth transitions between the active state and P1 without entering P2. Therefore, **Eq**. (**S6**) captures all possible ways of RNAp visiting P1 (and not P2 or P3) before taking one step forward.

Similar expressions can be derived for 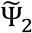 and 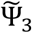, which are simplified assuming that P2 and P3 are visited at most once per step. This assumption is justified since the probability of entering P2 is estimated to be as low as ∼0.006 in the reference data set. Within this limit we obtain

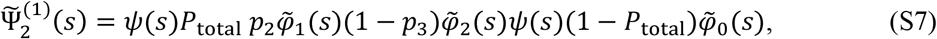

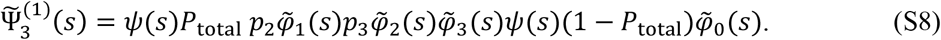

The term *ψ*(*s*) appears twice in **Eqs**. (**S7**) and (**S8**), accounting for back-and-forth transitions between the active state and P1 before and after entering P2, respectively.

Considering that the pause lifetimes, as estimated from the data, are well separated and much slower than the effective elongation rate (Table S2), each term in **Eq**. (**S1**) can be well approximated with a single exponential,

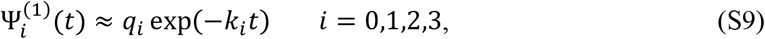

where *q*_*i*_ and *k*_*i*_ are the associated occurrence probabilities and effective rates respectively, and are related to the Laplace transform of 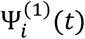 via the following equations:

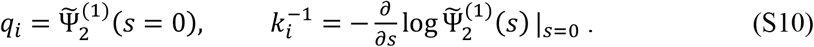

Considering that the Laplace transformation of an exponential function *φ*(*t*) = *k* exp(−*kt*) is given by 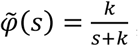, from **Eqs**. (**S5-8**) we obtain

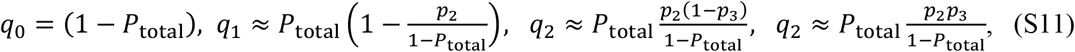

and

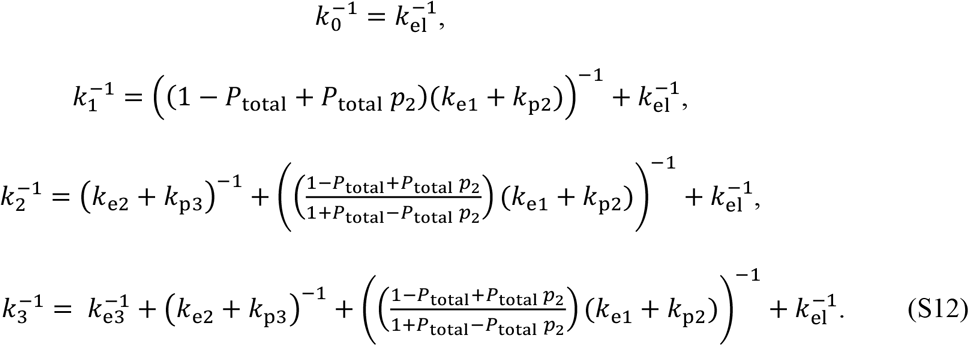

In obtaining the occurrence probabilities given in **Eq**. (**S11**) all terms of the order (*P*_total_*p* _2_)^2^ or greater have been ignored, in accordance with the assumption that multiple entrance to P2 is negligibly rare. This preserves the normalization of Ψ^(1)^(*t*), i.e. 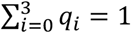. Replacing **Eq**. (**S9**) in **Eq**. (**S1**) yields

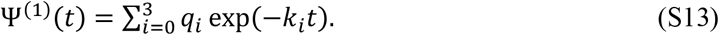

**Eq**. (**S13**) can be interpreted as RNAp exhibiting four distinct stepping rates, where at each step the rate *k*_*i*_ is chosen with the probability *q*_*i*_. While *k*_1_ = k_el_ corresponds to the pause-free elongation rate, each of the other three stepping rates are dominated by the exit rate from one of pause states in the model, as described in **Eq**. (**S12**). Considering that *k*_*i*_ s are sorted in the decreasing order (i.e. *k*_*i*_ > *k*_*i*+1_), this interpretation allows for the generalization of **Eq**. (**S1**) in the case where the width of the DWT-window, *N*, is larger than one nucleotide. One can thus describe

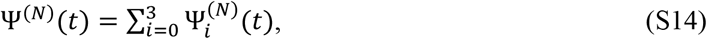

with 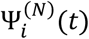 representing the conditional DWT distribution, where every step within the DWT-window is restricted to obtain a rate equal or greater than *k*_*i*_. For 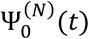, which corresponds to the pause-free elongation with the rate *k*_0_ = *k*_el_, we have

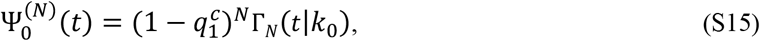

where 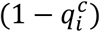 is the probability of not entering the pause branch, 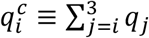, and the gamma function Γ_*n*_(*t*|*k*) represents the DWT distribution resulting from *n* successive steps with the rate *k*,

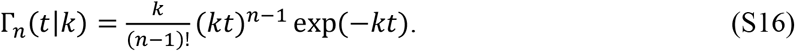

For *i* ≥ 1, 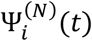 can be decomposed into *N* separate terms with

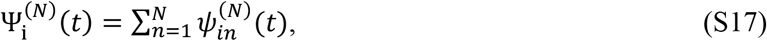

where 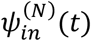 is conditioned to include exactly *n* pauses with the rate *k*_i_, while all the other steps in the DWT-window are taken at a rate much greater than *k*_i_. The fact that the stepping rates, as inferred from our data, are well separated, allows for a mean-field approximation for 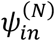 of the form

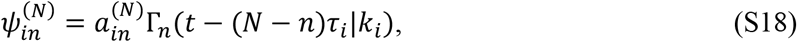

where 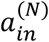 is the probability of RNAp taking *n* steps with the rake *k*_*i*_ and *N* − *n* steps with a rate faster than *k*_*i*_,

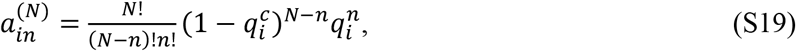

and we have included *τ*_*i*_ as the average DWT per step, given that the stepping rate is greater than *k*_*i*_,

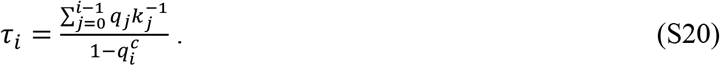

The gamma function Γ_*n*_(*t* − (*N* − *n*)*τ*_*i*_|*k*_*i*_) in **Eq**. (**S18**) accounts for the occurrence of *n* pauses with the lifetime 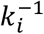. The shift (*N* − *n*)*τ*_*i*_ in the argument of the function reflects the mean-filed assumption that each of the remaining (*N* − *n*) steps contribute to the total DWT by a constant average time *τ*_*i*_. We assume 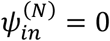 for *t* < (*N* − *n*)*τ*_*i*_.

Taken together, we obtain the following approximate analytical expression for the theoretical DWT distribution,

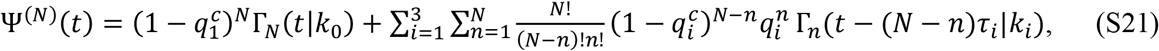

where *k*_*i*_s and *q*_*i*_s are related to the model parameters *via* **Eqs**. (**S11, 12**). We note that **Eq**. (**S21**) is reduced to (**S13**) for *N* = 1.

### M6: Estimation of the model parameters and number of kinetic states

We estimated the model parameters *via* a maximum likelihood fit to the reference data set (7.5 *pN* AF, 1 mM NTPs). Due to the existence of empirical noise, as well as the artifacts induced by the filtering process, **Eq**. (**S21**) cannot be directly used for fitting the data. While for pauses longer than the filtering time the noise artifacts can be suppressed by choosing a wide enough window (in this case 10 nt), the fast pause-free elongation rate, which is represented by the elongation peak in the DWT distribution, is significantly affected. In this case the elongation peak cannot be accurately described by a gamma function. We therefore kept the second term in **Eq**. (**S21**) to model the contribution from the pauses, but replaced the gamma function Γ_*N*_ (*t*|*k*_0_) by a log-normal function which better captures the elongation peak,

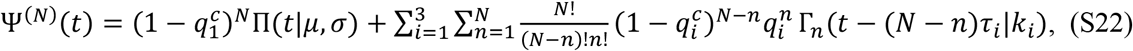

where

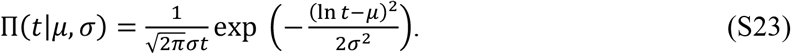

We fitted **Eq**. (**S22**) to the reference data, treating *μ* and *σ* as independent fit parameters. This introduces a challenge in estimating the effective elongation rate since an analytical formula which relates *μ* and *σ* to *k*_el_, and accurately accounts for the noise and filtering artifacts, is not known. We therefore kept *k*_*el*_ fixed at an initial guess and estimated all pause exit rates and probabilities (i.e. {*k*_*i*_. *q*_*i*_|*i* = 1.2.3}) by fitting **Eq**. (**S22**) to the reference data. The fit was performed by numerically maximizing the coarse-grained log-likelihood function defined as

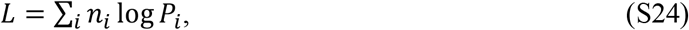

where *n*_*i*_ is the number of DWTs in the *i*_th_ bin of the empirical DWT distribution, and *P*_*i*_ is the model prediction calculated for that bin. The standard simulated annealing algorithm implemented in MATLAB was used in the optimization.

We then refined our initial guess for the elongation rate by performing a second numerical maximum likelihood fit, taking into account the empirical noise and filtering effects. We note that the elongation rate implicitly enters **Eq**. (**S22**) through the parameters *τ*_*i*_s. Our estimations for pause exit rates and probabilities can therefore be potentially affected by our initial guess for the elongation rate. We estimated *τ*_3_ for the reference data set by measuring the average pause-free velocity, excluding all pauses which are longer than the pause P3 lifetime of ∼100 s. This yielded *τ*_3_∼0.1 s, which also serves as an upper bound for *τ*_1_ and *τ*_2_. Therefore, for *N* = 10 we have *Nτ*_*i*_ ≤ 1 s, which is negligible compared to the estimated pause lifetimes of ∼5 s and ∼100 s for P2 and P3, respectively. This implies that that we can ignore the terms (*N* − *n*)*τ*_*i*_ for *i* = 2.3 in the argument of the gamma functions in **Eq**. (**S22**). As a result, we concluded that our estimations for *k*_2_, *k*_3_, *q*_2_ and *q*_3_ are not significantly affected by our choice of *k*_el_, as long as it does not lie far from the correct value. This argument does not necessary hold for the parameters associated with the pause P1, for which we have estimated a lifetime <1 s. Nevertheless, we expect that the estimation for *k*_1_ remains independent of *k*_el_, as it is dictated by the exponential decay in the tail of the DWT distribution at short time scales (i.e. between 1 - 4 s).

In contrast, by simulating DWT distributions from noisy traces (see below), we noticed that the estimation for *q*_1_ is indeed sensitive to the chosen value for *k*_el_. Based on this finding, to refine our initial guess for the elongation rate we only used *k*_el_ and *q*_1_ as free parameters in the numerical maximum likelihood fit, keeping *k*_1.2.3_ and *q*_2.3_ at their estimated values resulting from fitting **Eq**. (**S22**) to the data. The fit was performed *via* grid optimization. For each grid point, an ensemble of trajectories was computationally generated using the Gillespie algorithm.(66) To achieve an accurate estimation of the effective elongation rate, we included experimental noise. As the empirical noise turned out to be temporally correlated, we employed an ensemble of RNAp pauses exceeding 100 s as the noise source. After adding this noise to the computationally generated trajectories, the trajectories were filtered with a moving average filter of 1 Hz, as in the preprocessing or experimentally RNAp trajectories. The DWT distribution was then constructed from these computationally generated noisy trajectories. This allows for the calculation of the coarse-grained log-likelihood given by **Eq**. (**S24**), where *n*_*i*_ are the number of DWTs in the *i*_th_ bin of the empirical DWT distribution, and *P*_*i*_ is the simulated DWT distribution calculated for that bin. The gridpoints with the largest log-likelihood score yields an estimation for the optimal values of *k*_el_ and *q*_1_.

Having estimated the complete set of {*k*_*i*_. *q*_*i*_|*i* = 0.1.2.3}, we calculated all the model parameters by inverting **Eqs**. (**S11, S12**). Defining 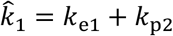 and 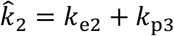, followed by

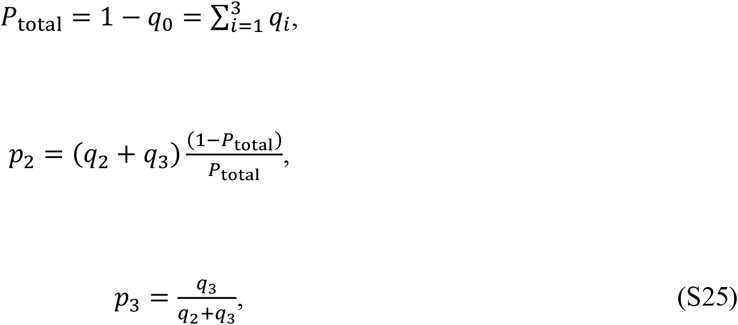

and

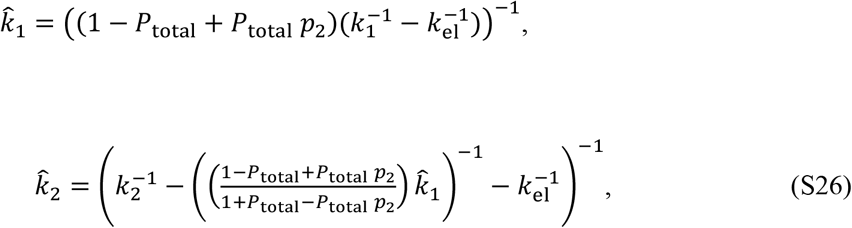

which in combination with **Eqs**. (**S3**) and (**S12**) followed by

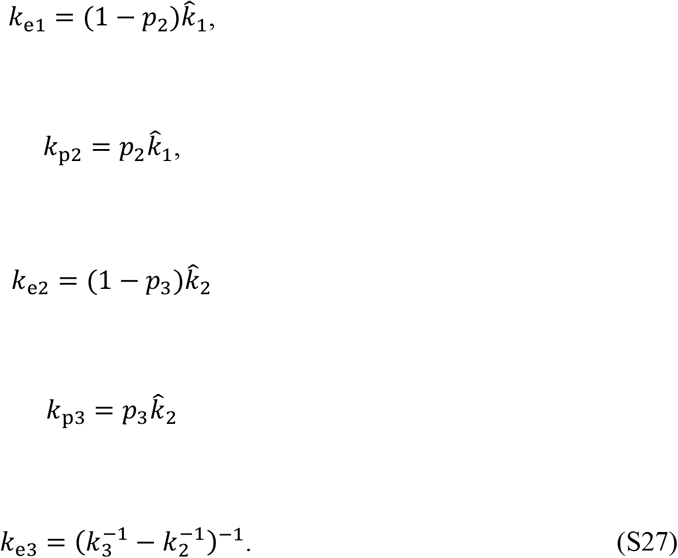

This completes the parameter estimation procedure. To estimate the 1-σ confidence intervals for the model parameters, the data was bootstrapped 100 times and the entire procedure was repeated for each bootstrapped sample.

**Eq**. (**S22**) can be generalized to describe a generic model, with an arbitrary number of *M* exponential pauses, by simply setting the upper bound of the first sum in **Eq**. (**S22**) to *M* instead of 3. This allowed us to repeat the fit for models with a different number of pause states (e.g. 2-6) and use the Bayesian Information Criteria (BIC) to confirm the existence of three distinct pause states in our data (24, 28).

### M7: Validation of the ML fit to the reference data set

The reference parameters (**Fig. 2B, C**) were obtained by fitting the model to the empirical DWT distribution using a DWT-window of 10 nt. To validate that the choice of the DWT-window does not affect the analysis, the reference parameters were fed to the Gillespie algorithm to computationally generate an ensemble of transcription trajectories as described in **M6**. The DWT distributions with the window sizes of 4 and 20 nt were then constructed from the computationally generated trajectories and compared to the empirical distributions **Fig S2B, C**.

To confirm that the value obtained for the effective elongation rate *k*_el_ can be accurately determined by the DWT analysis, it was reevaluated by fitting the pause-free velocity distribution as is commonly used in literature (11, 21). The local velocity of each RNAp trajectory was measured in successive time windows of 1 s, after removing pauses >100 s. These measured velocities were then used to construct the pause-free velocity distribution. The model was then fitted to the distribution by 1-D grid optimization with the effective elongation rate *k*_*el*_ as the only free fit parameters (Fig. **S2 E, F**); all other model parameters are kept fixed at their reference values (**Fig. 2B**,**C**). At each grid point an ensemble of trajectories were computationally generated and the pause-free velocity distribution was constructed. The coarse-grained log-likelihood score was then calculated at each grid point according to **Eq**. (**S24**), where *n*_*i*_ is the number of velocities in the *i*_th_ bin of the empirical pause-free velocity distribution, and *P*_*i*_ is the simulated distribution calculated for that bin. The data was bootstrapped and refitted 100 times to estimate the 1-σ confidence interval for the effective elongation rate *k*_el_.

To evaluate the variations in the elongation rate *k*_el_ and pause probability *P*_total_, the model was fitted to each individual trajectory in the reference data set. This was performed by a 2-D grid optimization with *k*_el_ and *P*_total_ as free fit parameters; all other model parameters were kept fixed at their reference values (**Fig. 2B**,**C**). For each grid point, the coarse-grained log-likelihood score (**Eq**. (**S24**)) was calculated using simulated noisy elongation trajectories as described in M6. The optimal values for *k*_el_ and *P*_total_, corresponding to the largest score, were then determined, and it was shown that the population-average estimations are consistent with the results from fitting the combined distribution (**Fig. S3D**)

### M8: Characterizing the change in model parameters at modulated empirical conditions

To fit the empirical DWT distributions measured at low NTP concentrations (i.e. 500 µM, 100 µM, 10 µM, and 1 µM), a 2-D grid optimization of the effective elongation rate *k*_el_ and pause probability *P*_total_ was performed by numerically maximizing the coarse-grained log-likelihood score as described in M7. Each RNAp trajectory was fitted individually and the population-averaged values for *k*_*el*_ and *P*_*total*_ were determined. The uncertainty in the fit outcome was characterized by calculating the standard error of the mean (**see Fig. 3**).

To characterize the force dependency of the P2 pause entry rate, *k*_p2_, the model was fitted to the DWT distribution measured at OF 12.5 pN, 1 mM NTPs. The fit was performed in two steps: first, for 11 equally-spaced values of *δ* in the interval [0,1] the best fit to the data was found using a 2-D grid optimization as described above for low-NTP fits. Each fit was performed using *k*_el_ and *P*_total_ as the only free fit parameters; *k*_p2_ was adjusted according to equation **Eq**. (**3**) and all the remaining model parameters were kept fixed at their reference values. The log-likelihood score corresponding to the best two-parameter fit was then calculated for each value of *δ*, and the optimal value of *δ* = 0.7 corresponding to the largest score was determined. To estimate the uncertainty in *δ*, the log-likelihood scores were exponentiated and normalized to obtain the probability distribution of *δ*, from which the 1-σ confidence interval was then calculated (**see Fig. 4**).

For all other externally applied forces (i.e. F= −10, −9, −7.5, −5, 5, 10, 12.5 pN) *k*_p2_ was adjusted according to **Eq**. (**3**) using *δ* = 0.7, and the model was fitted to the empirical DWT distribution using *k*_el_ and *P*_total_ as the only free fit parameters, as described above (see **Figs. 4, S5 B**,**C**).

The DWT distribution measured at 9 pN OF served as an individual control for the experiments in presence of GreA or GreB. The fit was performed in two steps. First, only *k*_e2_ and *k*_p3_ were considered as free fit parameters, while all other model parameters were fixed according to the best fit to the control data. The optimal values of *k*_e2_ and *k*_p3_ were determined by maximizing the coarse-grained log-likelihood score on a 2-D grid, calculated by simulating noisy elongation trajectories as described in M6 and M7. Then, a second maximum likelihood fit was performed on a 1-D grid with *k*_e3_ as the only free fit parameter. Due to low statistics, the fit was performed on the combined distribution and the data was bootstrapped and refitted 100 times to estimate the 1-σ confidence intervals for the fit outcome (see **Fig. 5A, B, C, D**).

### M9: The dependence of pause probability and effective elongation rate on NTP concentration and external force

The total pause probability, *P*_total_, is affected by the sub-processes involved the active pathway. As depicted in **Fig. S4B**, these sub-processes include the transition between pre- and post-translocated states (denoted by 0 and 1, respectively) with the forward rate *k*_*F*_ and the backward rate *k*_*B*_, the transition to the NTP-bound state (denoted by 2) with the rate *k*_*N*_, NTP-dissociation with the rate *k*_*D*_ and finally the NTP catalysis with the rate *k*_*C*_. The catalytic reaction completes the elongation cycle and brings the RNAp back to the next pre-translocated state (denoted by 3), with a new nucleotide added to the RNA chain. Defining *p*_*ij*_ as the transition probability between two neighboring sub-states *i* and *j* = *i* ± 1, the probability to complete the elongation cycle without entering the pause branch can be written as

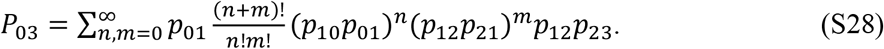

**Eq**. (**S28**) accounts for all possible paths from 0 to 3 which do not include entering the pause branch: each path begins with a transition from 0 to 1 followed by *n* back-and-forth transitions between 1 and 0 and *m*

back-and-forth transitions between 1 and 2, all ending at 1. The factor 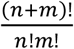 accounts for all possible permutations for such transitions. A final transition from 1 to 2 followed by a transition from 2 to 3 then completes the catalytic pathway. Using the binomial theorem, **Eq**. (**S28**) can be simplified as

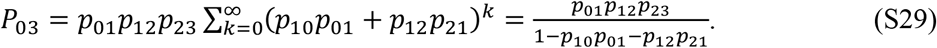

By definition

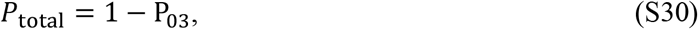

and the probabilities *p*_*ij*_ are related to the transition rates as follows

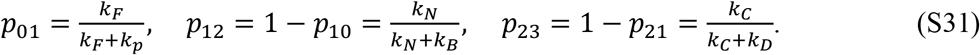

Considering that *k*_*N*_ is proportional to the NTP concentration, 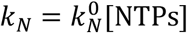, **Eq**. (**S30**), together with (**S29**) and (**S31**), yields

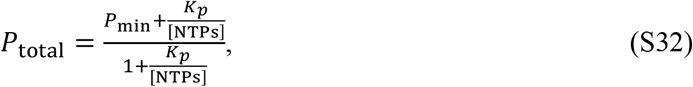

where

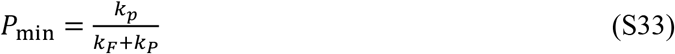

is the lowest value of *P*_total_ corresponding to a saturating NTPs concentration, and

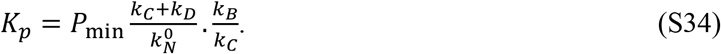

The transition rates between the pre- and post-translocated states depend on the external force *F* as

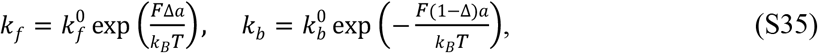

where *a* is the average step size of RNAp, 0 ≤ Δ ≤ 1 identifies the position of the energy barrier against RNAp transition from the pre-translocated to the post-translocated state, and 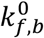 corresponds to zero force. Assuming that the entry rate to the pause branch, *k*_p1_, does not depend on force, using **Eqs**. (**S33-35**) we obtain

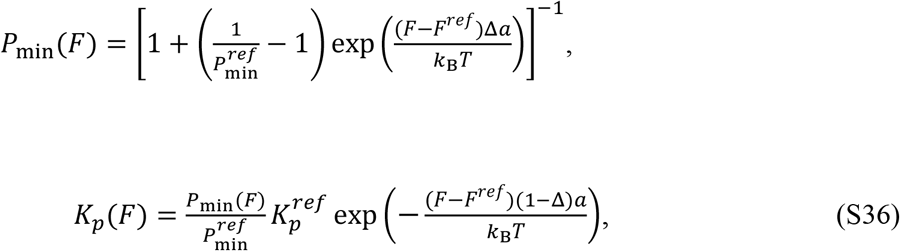

where 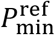 and 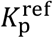 correspond to a reference force *F*^ref^.

An analytical expression can be obtained for the effective elongation rate. We denote the first-passage time distribution of completing one elongation cycle without pausing (i.e. transiting from state 0 to3 in **Fig. S4B**) by Ψ_03_ (*t*), and its Laplace transformation by 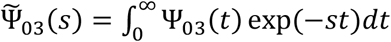. Similar to **Eq**. (**S29**) one can write

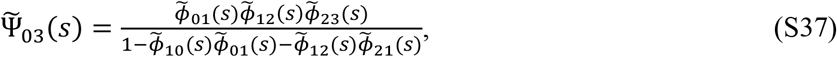

where 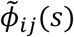 is the Laplace transformation of the first-passage time distribution for a transition from state *i* to a neighboring state *j*,

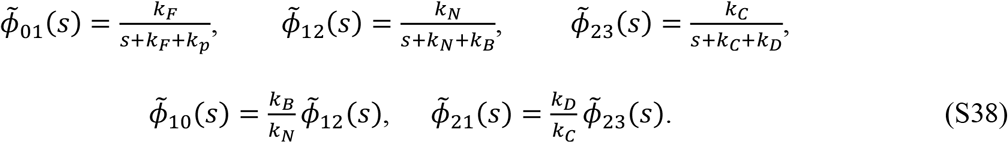

The elongation rate, k_el_, can then be calculated as

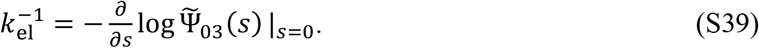

Using **Eq**. (**S38**) and defining

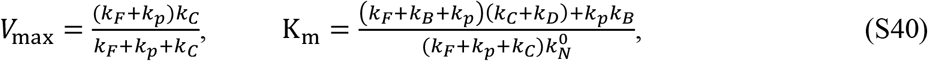

we obtain

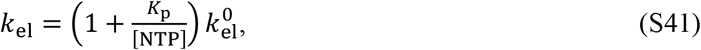

where *K*_p_ is given by **Eq**. (**S34**) and 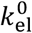 is the elongation rate at *k*_*p*_ = 0 which follows the Michaelis-Menten kinetics

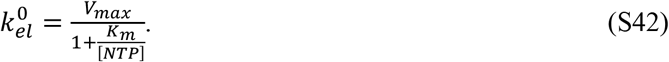

### M10: Parameter estimation for the diffusive backtracking model

To model diffusive backtracking, we modified equation **Eq**. (**S22**) to include one exponential pause with the rate *k*_1_ and probability *q*_1_, as well as a diffusive backtracking pause with the probability *q*_2_, the forward rate *k*_*f*_, and the backward rate *k*_*b*_. Assuming *Nq*_2_ ≪ 1, which is justified for *N* = 10 and *q*_2_∼0.006, we obtain

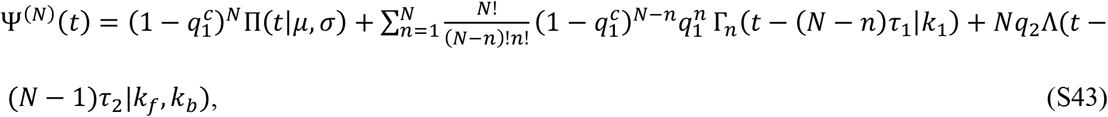

where 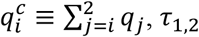, *τ*_1,2_ are given by **Eq**. (**S20**), and Λ(*t*|*k*_*f*_. *k*_*b*_) is given by

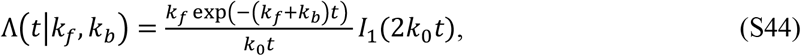

with 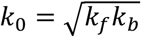, and *I*_1_(*x*) is the degree one modified Bessel function of the first kind (19).

The diffusive backtracking model was fitted to the reference data using a similar procedure as described in M6. First a maximum likelihood fit was performed using **Eq**. (**S43**), with *k*_*el*_ kept fixed at an initial guess. A second maximum likelihood fit *via* 2-D grid optimization was then performed using *q*_1_ and *k*_el_ as the only free parameters. While the forward and backward rates *k*_*f*_, *k*_*b*_ were directly estimated from the fit, estimating *k*_el_, *k*_1_, and *q*_1,2_ allows for adjusting **Eqs**. (**S25-27**) to calculate *P*_total_, *k*_e1_, and *kp*_BT_ as follows (Figure S6, rightmost panel):

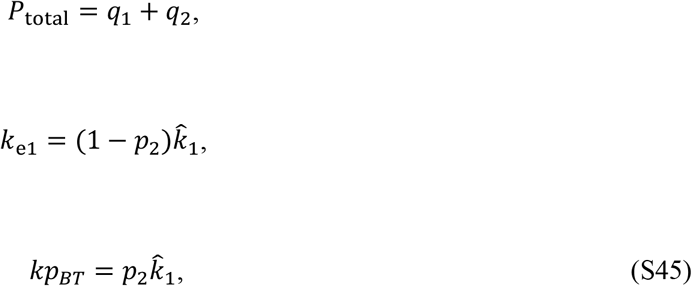

where 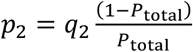 and 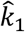 is given by **Eq**. (**S26**). Using the estimated model parameters, the DWT distribution was simulated with limited diffusive backtracking, were the maximum backtrack depth was varied between 3 − 14 nt.

The model was tested on the data measured at OF 12.5 pN and 1 mM NTPs, assuming that the rates associated with backtracking depend on force *via* the following equations:

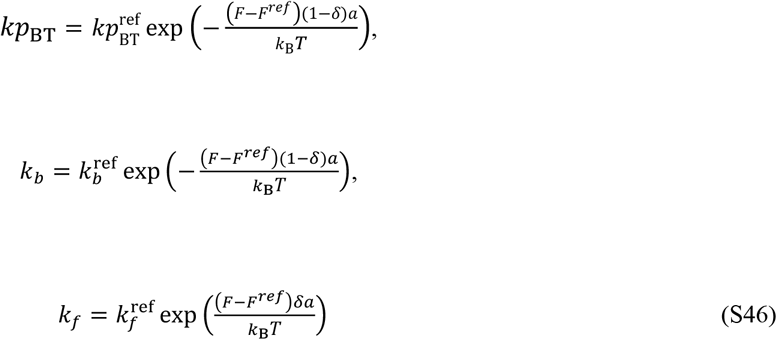

where *F* = −12.5 pN, *a* ≈ 0.37 nm, and the superscript “*ref*” refers to the value of the parameter estimated for the reference data at *F*^ref^ = 7.5 pN. An ensemble of noisy traces was then simulated as described in M6 to construct DWT distributions and calculate the average RNAp processivities for different values of *δ* and maximum backtrack depth. In all simulations, *k*_el_ and *p*_total_ were fixed according to Table S2. The best fit of the diffusive backtracking model to the 12.5 *pN* OF data was find by calculating the coarse-grained log-likelihood score using **Eq**. (**S24**) for each simulation.

### M9: Statistical analysis

Statistical significance tests were performed by using unpaired, two tailed t-tests and one-way analyses of variance (ANOVA) with Tukey post hoc tests. The levels of significance are denoted in the figure captions. All data plot values were shown as mean average ±SD.

## Notes

### Competing Interest Statement

The authors have declared no competing interest.

