## Supplementary information for "Accounting for RNA polymerase heterogeneity reveals state switching and two distinct long-lived backtrack states escaping through cleavage"

**This PDF file includes:**

Figures S1 to S6

Tables S1 to S3

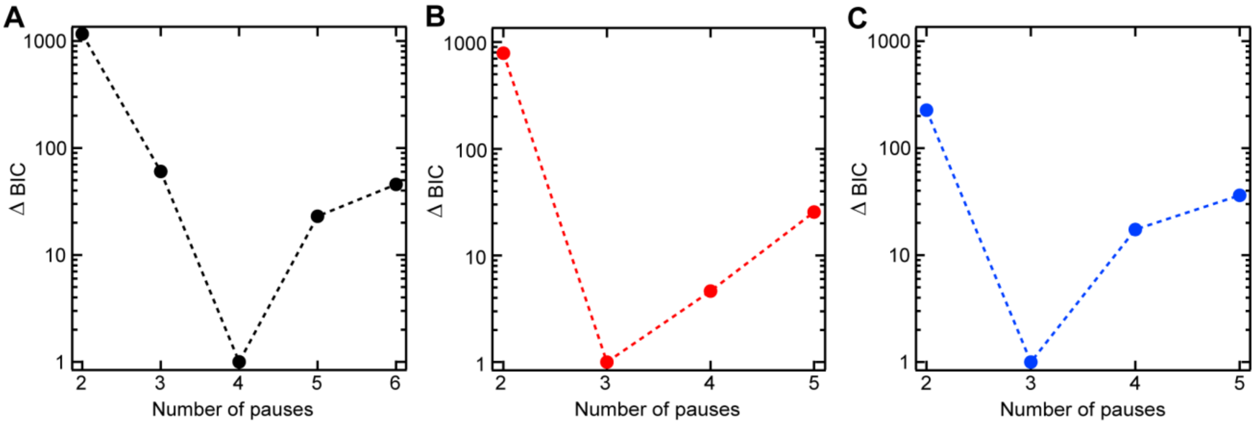


**Figure S1: Bayesian information criterion (BIC) analysis of pausing dynamics.** (**A**) All RNAp trajectories in the reference data set (7.5 pN AF, 1 mM NTPs) were used to construct a single combined DWT distribution. Models with a different number of single-rate pause states were fitted to the DWT distribution, and the corresponding BIC values were computed (Methods M5 and M6) (Press *et al*., 2007; Schwarz, 1978). The minimal BIC value reveals the optimal estimation for the number of single-rate pause states. Panels (**B**) and (**C**) depict the BIC estimation for the dominant and minority populations shown in Figure 1B, respectively, separated by our selection protocol (Methods M4). **Related to Figures 1 and 2**.

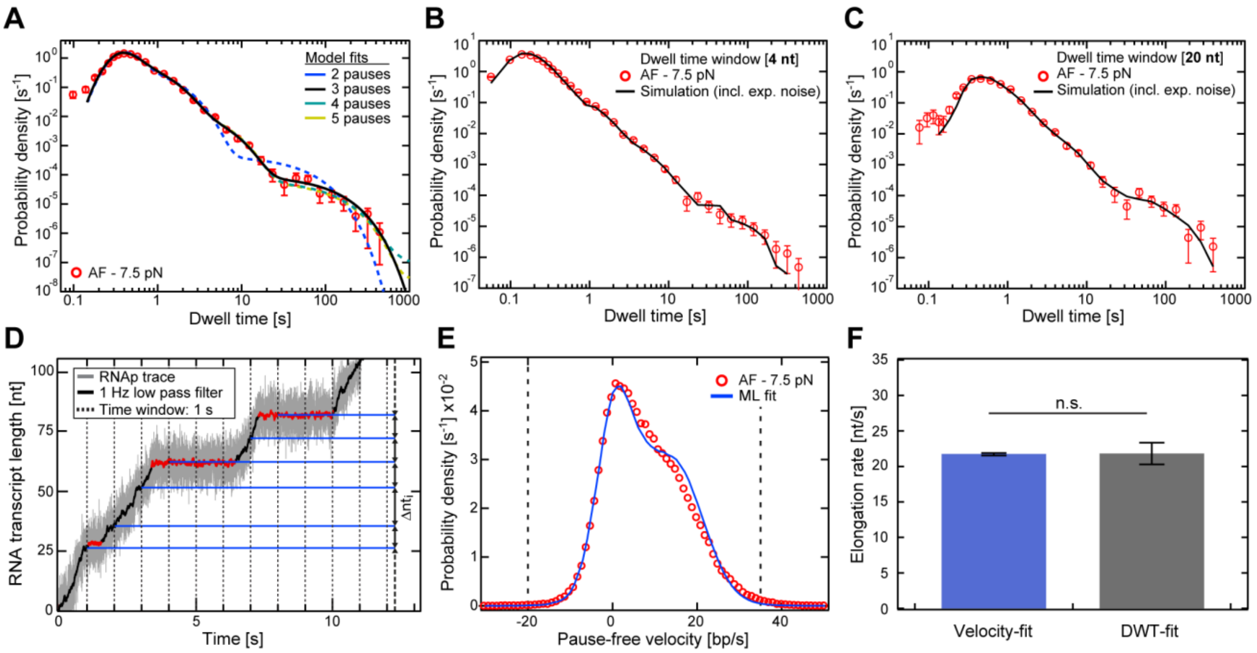


**Figure S2: DWT and velocity distribution analysis of RNAp transcription dynamics.** (**A**) ML fitting of multiple single exponential pauses states (2 - 5 pause states; colored lines) to empirical DWT distribution (red circles; condition: 7.5 pN AF, 1 mM NTPs). (**B**, **C**) Superimposition of empirical data (red circles) and the simulated DWT distributions as predicted by our model (Figure 2, rightmost panel) using two DWT-window sizes. (**D**) An example of a measured RNAp trajectory before (gray) and after (black) filtering to 1 Hz. To calculate the local "pause free" velocities of RNAp, the portions of the trajectory that correspond to DWTs longer than 100 s (red) are removed and the distance traversed by the RNAp (blue horizontal lines) in the successive widows of 1 s (grey vertical dashed lines) is measured. (**E**) The empirical distribution (red circles) of the RNAp pause-free velocity (as defined in D) constructed from pooled RNAp trajectories measured at 7.5 pN AF, 1 mM NTPs. The blue curve represents the best model fit where the effective elongation rate $k_{el}$ is the only free parameter while all other parameters are kept constant as in Figure 2B,C. (**F**) The resulting values for $k_{el}$ determined by fitting the pause-free velocity distribution (blue) compared to the value obtained by fitting to our DWT distribution (gray; Figure 2A,B). **Related to Figure 2**.

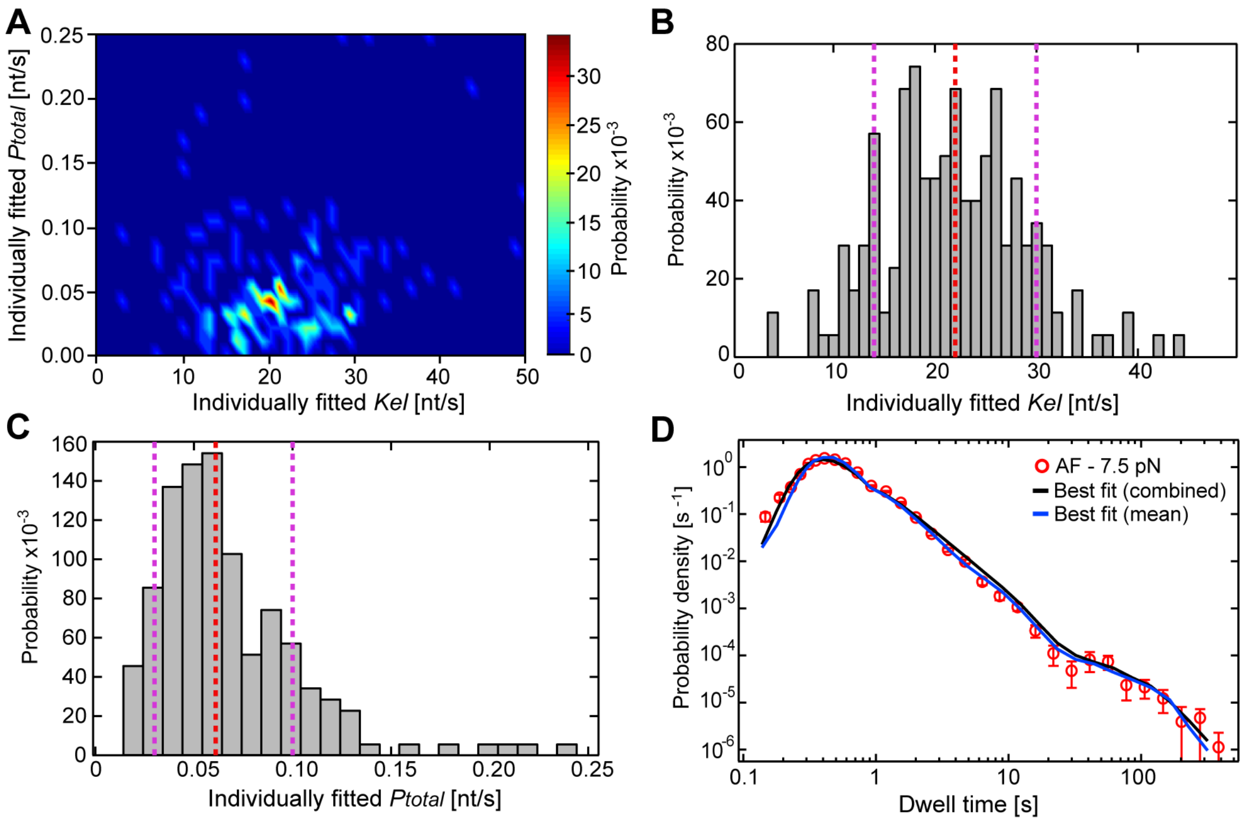


**Figure S3: Estimation of effective elongation rate and total pause probability in individual RNAp trajectories.** (**A**) Two-dimensional distribution of the effective elongation rate $k_{\mathrm{el}}$ and the total pause probability $P_{\mathrm{total}}$ obtained by fitting the model to individual transcription trajectories (Methods M8). Pearson correlation coefficient ($r$ = 0.16854) indicates no significant correlation between $k_{\mathrm{el}}$ and $P_{\mathrm{total}}$. (**B**), **(C)** One-dimensional distributions of $k_{\mathrm{el}}$ and $P_{\mathrm{total}}$, respectively, from the data used to construct (A). The red dashed lines mark the average of the distributions, and the purple dashed lines identify the 1-σ confidence interval. **(D)** The best fit (black solid line) to the combined DWT distribution measured at 7.5 pN AF, 1 mM NTPs (red circles). The best fit was obtained by estimating $k_{el}$ and $P_{\mathrm{total}}$ for individual transcription trajectories (see (B) and (C)), and simulating the corresponding individual DWT distributions. The weighted average of the simulated distributions was then calculated, where each distribution was weighted with the number of DWTs that exist in each measured trajectory. The blue solid line depicts the simulated DWT distribution with the averaged population values of $k_{\mathrm{el}}$ and $P_{\mathrm{total}}$ (red dashed lines in (B) and (C)). **Related to Figure 2**.

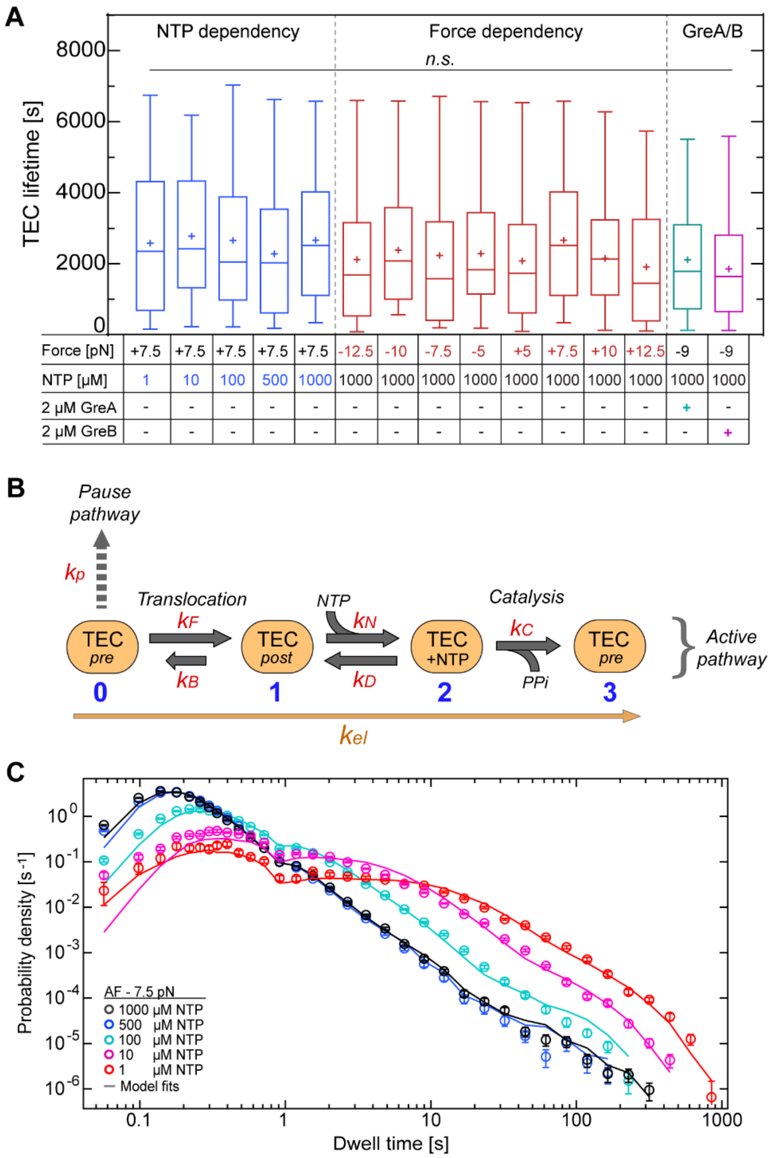


| Force [pN] | +7.5 | +7.5 | +7.5 | +7.5 | +7.5 | -12.5 | -10 | -7.5 | -5 | +5 | +7.5 | +10 | +12.5 | -9 | -9 |
|---|---|---|---|---|---|---|---|---|---|---|---|---|---|---|---|
| NTP [µM] | 1 | 10 | 100 | 500 | 1000 | 1000 | 1000 | 1000 | 1000 | 1000 | 1000 | 1000 | 1000 | 1000 | 1000 |
| 2 µM GreA | - | - | - | - | - | - | - | - | - | - | - | - | - | + | - |
| 2 µM GreB | - | - | - | - | - | - | - | - | - | - | - | - | - | - | + |

**Figure S4: Measured TEC lifetimes and processes of the catalytic pathway.** (**A**) Distribution of TEC lifetimes observed at different NTP concentrations (blue), different range of applied forces (red), and in presence of the transcription cleavage factors GreA (cyan) and GreB (purple). Measured lifetimes were subject to statistical analysis using one-way ANOVA with comparative Tukey post-hoc test. The different lifetimes exhibit no statistical difference (*n.s.* = *non-significant*). (**B**) Schematic of the sub-processes involved in the catalytic pathway. From left to right: reversible transition from a pre- to a post-translocated state, reversible NTP-binding, and irreversible NTP catalysis. (**C**) Superimposed DWT distributions (circles) of combined elongation trajectories at different NTP concentrations. The lines represent the model predictions whereby $k_{\mathrm{el}}$ was calculated using equations **Eq. 1** and **Eq. 2**, with $V_{\mathrm{max}} = 21.8\ \mathrm{nt/s}$, $K_{\mathrm{p}} = 8.6\ \mu\mathrm{M}$, and $\mathrm{K_m} = 100\ \mu\mathrm{M}$; all other model parameters were chosen according to the values in Table S1. **Related to Figures 3, 4, and 5.**

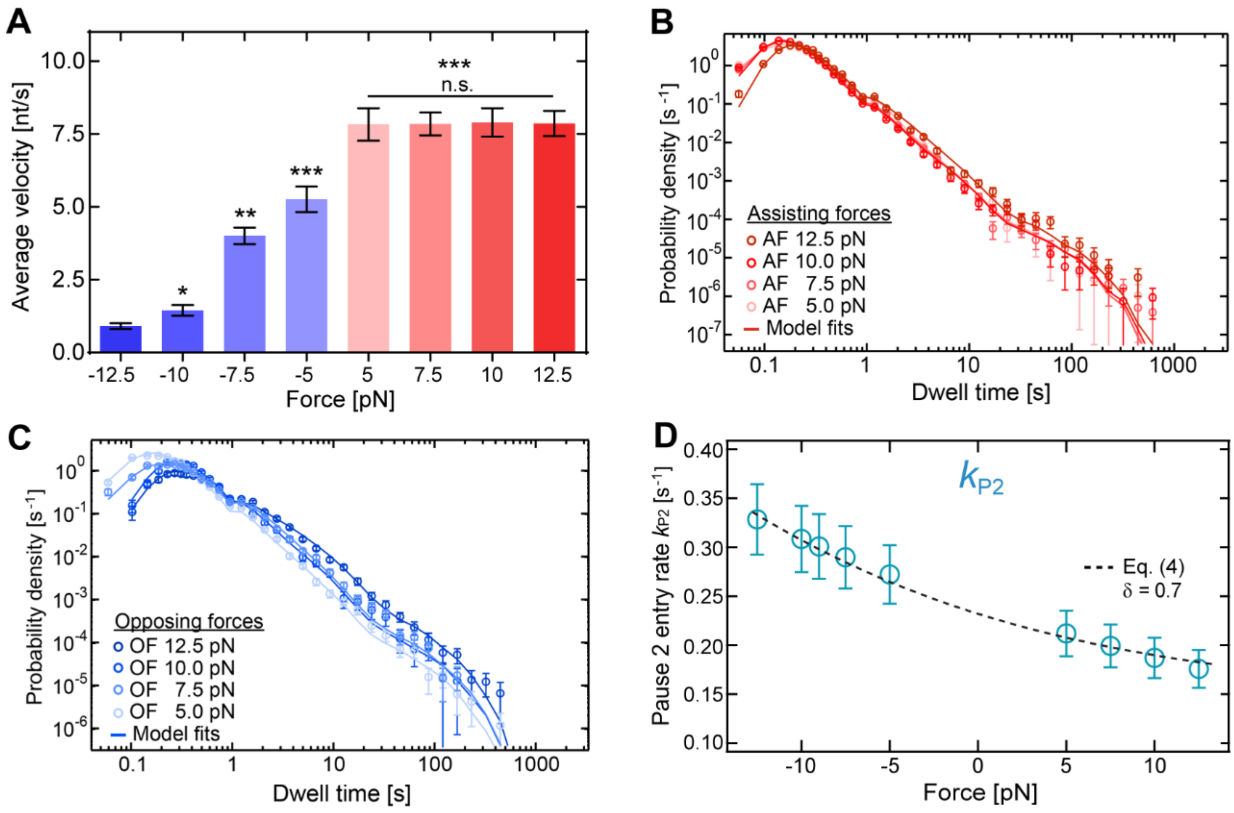


**Figure S5: Impact of different applied forces on average transcription velocity and dynamics.** (**A**) Average end-to-end velocities (±SEM) of measured elongation traces in function of the applied external force. The minus sign denotes opposing forces. Data were subject to statistical analysis using one-way ANOVA, with comparative Tukey post-hoc test (significance level p: * = 0.05, ** = 0.01, *** = 0.001, *n.s. = non-significant*). (**B**) Resulting best model fits to the data obtained at different magnitudes of applied assisting forces. (**C**) The same as B, but for a different magnitude of opposing forces. (**B,C**) Only the elongation rate $k_{el}$ and the pause entry probability $P_{total}$ were used as free fit parameters. The parameter $\delta$, which dictates the force dependency of $k_{P2}$, is fixed at its optimal value of 0.7 (Figure 4B). All other model parameters were fixed to their optimal values in Table S1. (**D**) $P_{total}$ probabilities from the fits to data sets at different applied forces (B) and (C). Error bars for (B, C) were determined by bootstrapping as described in the methods section. **Related to Figure 4.**

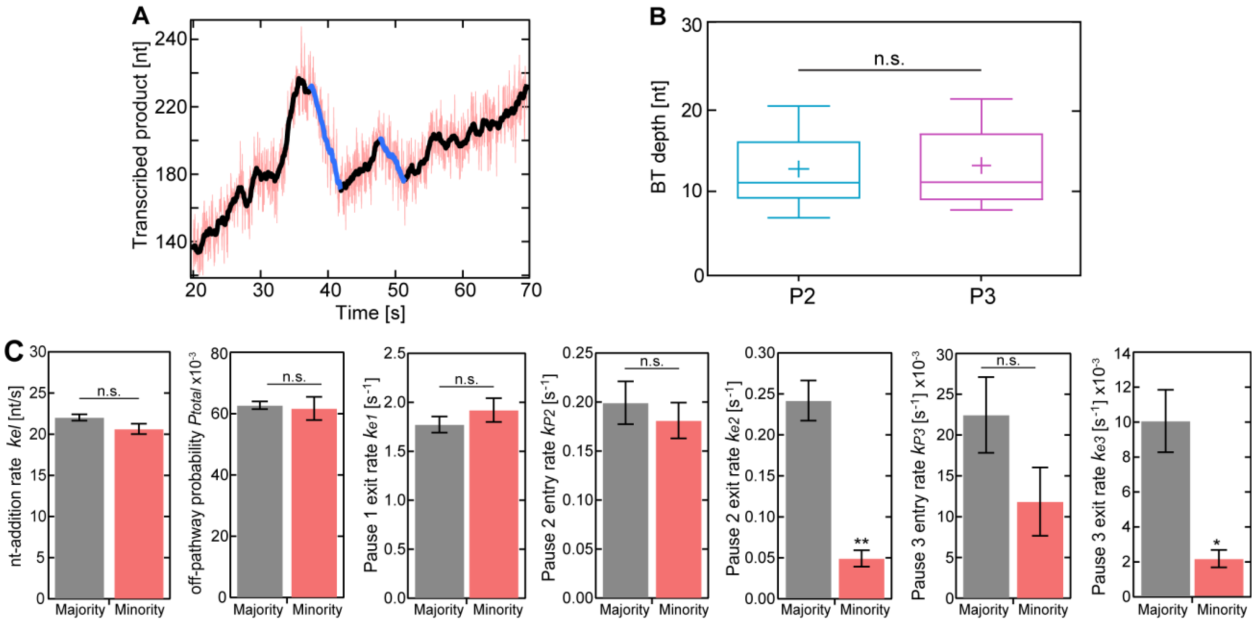


**Figure S6: Measured backtrack depths associated with P2 and P3 pause states**. (**A**) A magnified part of an example RNAp trajectory (red: raw data; black: 1 Hz filtered data) showing a directly observed backtrack by RNAp (blue). (**B**) Measured backtrack depths (in nt), separated by the average recovery times associated with the backtracked pause states P2 and P3. Whiskers represent 95% confidence intervals. (**C**) The optimal model parameters obtained by fitting the model to the minority (i.e. excluded) population (red), compared to the parameters corresponding to the majority (i.e. selected) population (gray). **Related to Figures 5 and 6.**

**Table S1: Single molecule experimental parameters and measurement statistics.** Table of experimental parameters, such as NTP concentrations, presence of GreA or GreB (2 µM), and magnitude and sign of the applied force. As statistical measure for the different experiments, the calculated dwell times and total length of synthesized RNA nucleotides from pooled RNAp trajectories are shown. Also depicted are the number (n) of RNAp trajectories analyzed for the majority (and minority) subpopulations. The dwell times are obtained using a dwell-time window of 4 nt with 1 Hz filtering.

| **[NTPs]** | 1000 µM | 500 µM | 100 µM | 10 µM | 1 µM | 1000 µM |
|---|---|---|---|---|---|---|
| **Force [pN]** | 7.5 | 7.5 | 7.5 | 7.5 | 7.5 | -12.5 |
| **GreA** | - | - | - | - | - | - |
| **GreB** | - | - | - | - | - | **-** |
| **Trajectories [n]** | 177 (19) | 138 (18) | 115 (20) | 67 (9) | 47 (11) | 45 (5) |
| **Dwell times** | **90320** | **55003** | **39748** | **28946** | **9334** | **8283** |
| **Nucleotides** | 361.3 kb | 220 kb | 159 kb | 115.8 kb | 37.3 kb | 33.1 kb |

| **[NTPs]** | 1000 µM | 1000 µM | 1000 µM | 1000 µM | 1000 µM | 1000 µM |
|---|---|---|---|---|---|---|
| **Trajectories [pN]** | -10 | -9 | -7.5 | -5 | 5 | 10 |
| **GreA** | - | - | - | - | - | - |
| **GreB** | **-** | **-** | **-** | **-** | **-** | **-** |
| **RNAp [n]** | 53 (11) | 37 (6) | 46 (5) | 71 (9) | 85 (20) | 171 (11) |
| **Dwell times** | **9830** | **21162** | **32448** | **21607** | **43030** | **76701** |
| **Nucleotides** | 39.3 kb | 84.6 kb | 129.8 kb | 86.4 kb | 172.1 kb | 306.8 kb |

| **[NTPs]** | 1000 µM | 1000 µM | 1000 µM |
|---|---|---|---|
| **Force [pN]** | 12.5 | -9 | -9 |
| **GreA** | - | + | - |
| **GreB** | **-** | **-** | + |
| **Trajectories [n]** | 145 (13) | 93 (10) | 128 (5) |
| **Dwell times** | **54787** | **23214** | **28976** |
| **Nucleotides** | 219.1 kb | 92.8 kb | 115.9 kb |

**Table S2: Kinetic model parameters (±SD) emerging from fits to measured RNAp data under different experimental conditions.** The reference values, corresponding to 7.5 pN AF and 1 mM NTPs, are denoted in blue. In all other cases only a subset of model parameters (denoted in red) were fitted, while the rest of the parameters (black) were kept fixed at their reference/control values.

| Force [pN] | NTPs [mM] | GreA or GreB | $k_{el}$ | $k_{e1}$ | $k_{e2}$ | $k_{e3}$ | $P_{total}$ | $k_{P2}$ | $k_{P3}$ |
|---|---|---|---|---|---|---|---|---|---|
| 12.5 | 1 | - | 22.86 ±0.63 | 1.77 ±0.08 | 0.241 ±0.024 | 0.0100 ± 0.0018 | 0.059 ±0.003 | 0.176 ±0.019 | 0.022 ±0.005 |
| 10 | 1 | - | 22.77 ±0.65 | 1.77 ±0.08 | 0.241 ±0.024 | 0.0100 ± 0.0018 | 0.065 ±0.005 | 0.187 ±0.021 | 0.022 ±0.005 |
| 7.5 | 1 | - | 22.04 ±0.39 | 1.77 ±0.08 | 0.241 ±0.024 | 0.0100 ± 0.0018 | 0.063 ±0.001 | 0.199 ±0.022 | 0.022 ±0.005 |
| 5 | 1 | - | 21.77 ±0.79 | 1.77 ±0.08 | 0.241 ±0.024 | 0.0100 ± 0.0018 | 0.037 ±0.003 | 0.212 ±0.023 | 0.022 ±0.005 |
| -5 | 1 | - | 16.58 ±0.78 | 1.77 ±0.08 | 0.241 ±0.024 | 0.0100 ± 0.0018 | 0.065 ±0.005 | 0.289 ±0.032 | 0.022 ±0.005 |
| -7.5 | 1 | - | 14.79 ±1.09 | 1.77 ±0.08 | 0.241 ±0.024 | 0.0100 ± 0.0018 | 0.168 ±0.007 | 0.272 ±0.029 | 0.022 ±0.005 |
| -9 | 1 | - | 23.34 ±1.69 | 1.77 ±0.08 | 0.241 ±0.024 | 0.0100 ± 0.0018 | 0.117 ±0.015 | 0.301 ±0.033 | 0.022 ±0.005 |
| -10 | 1 | - | 13.01 ±1.04 | 1.77 ±0.08 | 0.241 ±0.024 | 0.0100 ± 0.0018 | 0.166 ±0.011 | 0.308 ±0.033 | 0.022 ±0.005 |
| -12.5 | 1 | - | 16.95 ±1.67 | 1.77 ±0.08 | 0.241 ±0.024 | 0.0100 ± 0.0018 | 0.256 ±0.017 | 0.328 ±0.036 | 0.022 ±0.005 |
| 7.5 | 0.5 | - | 21.43 ±0.76 | 1.77 ±0.08 | 0.241 ±0.024 | 0.0100 ± 0.0018 | 0.057 ±0.003 | 0.199 ±0.022 | 0.022 ±0.005 |
| 7.5 | 0.1 | - | 20.32 ±0.91 | 1.77 ±0.08 | 0.241 ±0.024 | 0.0100 ± 0.0018 | 0.202 ±0.006 | 0.199 ±0.022 | 0.022 ±0.005 |
| 7.5 | 0.01 | - | 23.91 ±1.29 | 1.77 ±0.08 | 0.241 ±0.024 | 0.0100 ± 0.0018 | 0.529 ±0.015 | 0.199 ±0.022 | 0.022 ±0.005 |
| 7.5 | 0.001 | - | 17.05 ±1.11 | 1.77 ±0.08 | 0.241 ±0.024 | 0.0100 ± 0.0018 | 0.797 ±0.009 | 0.199 ±0.022 | 0.022 ±0.005 |
| -9 | 1 | 2 µM GreA | 23.34 ±1.69 | 1.77 ±0.08 | 0.251 ±0.003 | 0.0120 ± 0.0005 | 0.117 ±0.015 | 0.301 ±0.033 | 0.015 ±0.003 |
| -9 | 1 | 2 µM GreB | 23.34 ±1.69 | 1.77 ±0.08 | 0.460 ±0.035 | 0.0110 ±0.0005 | 0.117 ±0.015 | 0.301 ±0.033 | 0.0197 ±0.0001 |

**Table S3: Primer sequences used for the synthesis of linear DNA tether constructs containing T7A1 promoter.**

| Oligonucleotide | Sequence |
|---|---|
| 1 | 5'- GACCGAGATAGGGTTGAGTG |
| 2 | 5'- CCATCTTGGTCTCCCCTACGCTCTAGAACTAGTGGATCCCCC |
| 3 | 5'- CCATCTTGGTCTCCTAGGCGTCAGCCTGCGAAGCAGTGGC |
| 4 | 5'- CCATCTTGGTCTCCTGTCAACACCACTTTGCTCCGAGGTT |
| 5 | 5'- CCATCTTGGTCTCCGACAGCGCCATTCGCCATTCAGGCTG |
| 6 | 5'- CTTCTGCTTTCCTGATGCAAAAAC |
| 7 | 5'- CTGCGGTCTCGCCCGACCGGCTCCAGATTTATCAGC |
| 8 | 5'- CTGCGGTCTCGTCAAAAACTGGAACAACACTCAACCC |
| 9 | 5'- CTGCGGTCTCGTAGGAGGCGCCATTCGCCATTCAGG |
| 10 | 5'- CTGCGGTCTCGCCGGGTTGCAGCACTGGGGCCAGATG |
